# Artificial Intelligence Helps to Predict Recurrence and Mortality for Prostate Cancer Using Histology Images

**DOI:** 10.1101/2023.07.27.550781

**Authors:** Okyaz Eminaga, Fred Saad, Zhe Tian, Ulrich Wolffgang, Pierre I. Karakiewicz, Véronique Ouellet, Feryel Azzi, Tilmann Spieker, Burkhard M. Helmke, Markus Graefen, Xiaoyi Jiang, Lei Xing, Jorn H. Witt, Dominique Trudel, Sami-Ramzi Leyh-Bannurah

**Affiliations:** AI Vobis, Palo Alto, CA; Division of Urology, Department of Surgery, Centre Hospitalier de l’Université de Montréal, University of Montreal, Montréal, QC H2X 0A9; Cancer Prognostics and Health Outcomes Unit, University of Montreal Health Center, Montreal, QC, Canada; University of Münster, Münster, Germany; Centre de recherche du Centre Hospitalier de l’Université de Montréal (CRCHUM), 900 Saint-Denis, Montréal, QC H2X 0A9, Canada; Institut du cancer de Montréal, 900 Saint-Denis, Montréal, QC H2X 0A9, Canada; Department of Pathology and Cellular Biology, Université de Montréal, 2900 Boulevard Édouard-Montpetit, Montreal, QC H3T 1J4, Canada; Department of Pathology, Centre Hospitalier de l’Université de Montréal (CHUM), 1051 Sanguinet, Montreal, QC H2X 0C1, Canada; Institute of Pathology, St. Franziskus-Hospital, Muenster, Germany Department of Pathology, University of Muenster, Germany; Institute of Pathology, Elbe Klinikum Stade, Academic Teaching Hospital of The University Medical Center Hamburg-Eppendorf (UKE), Hamburg, Germany; Martini-Klinik Prostate Cancer Center, University Hospital Hamburg-Eppendorf, Hamburg, Germany; Department of Computer Science, University of Muenster, Muenster, Germany; Department of Radiation Oncology - Radiation Physics, Stanford University School of Medicine, Stanford, CA; Prostate Center Northwest, Department of Urology, Pediatric Urology and Uro-Oncology, St. Antonius-Hospital, Gronau, Germany

## Abstract

Besides grading, deep learning could improve expert consensus to predict prostate cancer (PCa) recurrence. We developed a novel PCa recurrence prediction system based on artificial intelligence (AI). We validated it using multi-institutional and international datasets comprising 2,647 PCa patients with at least a 10-year follow-up. Survival analyses were performed and goodness-of-fit of multivariate models was evaluated using partial likelihood ratio tests, Akaike’s test, or Bayesian information criteria to determine the superiority of our system over existing grading systems. Comprehensive survival analyses demonstrated the effectiveness of our AI- system in categorizing PCa into four distinct risk groups. The system was independent and superior to the existing five grade groups for malignancies. A high consensus level was observed among five blinded genitourinary pathology experts in ranking images according to our prediction system. Therefore, AI may help develop an accurate and clinically interpretable PCa recurrence prediction system, facilitating informed decision-making for PCa patients.

## Introduction

Prostate cancer (PCa) is one of the most prevalent malignant diseases in males and exhibits diverse cancer aggressiveness and prognosis^1^. When PCa is diagnosed, usually by biopsy, the pathological examination of cancer differentiation and dissemination status are key determinants for selecting appropriate treatments^2^. Currently, pathologists grade PCa malignancy based on the modified Gleason grading system, originally established in the 1960s^3^. The first version of the Gleason grading system was based on five tissue patterns (labeled 1–5) that identified different transformation conditions of prostatic tissues according to tissue architecture, growth, and glandular features^3,4^. This grading system produces a score that considers two identical or different patterns to grade PCa differentiation, and the order in which patterns are added differs according to tissue sampling (biopsy core vs. whole prostate)^3,4^. PCa grading was further refined after patterns 1 and 2 were mostly identified as benign with the identification of basal cells by immunohistochemistry, and some of those patterns 1 and 2 were reclassified as Gleason pattern 3 as well^5,6^. In 2016, Epstein *et al*. proposed a modified version of the Gleason grading system that included five grade groups (GGs) instead of nine different Gleason scores (such as 3 + 3, 4 + 3, and 5 + 3) to achieve a more concise prognostic stratification according to biochemical recurrence (BCR) rates^7^.

Despite strong prognostic capacities and continual revisions since its introduction^8^, GG reproducibility has remained limited because of interobserver variability in grading and quantification, leading to grade inconsistency even among expert pathologists, thus increasing the potential risk of treatment delay or suboptimal treatment choice^9,10^. Contemporary studies have highlighted the great potential of artificial intelligence (AI) in improving GG consistency and achieving accuracy comparable to expert levels^11–13^. However, these studies likely inherited the limitations of the current grading system as the histological ground truth is based on evaluations from a small group of expert pathologists, which is not necessarily reflective of the global pathology community (social and cognitive biases) or grading correctness^14^.

To bypass these reproducibility limitations, we applied AI to develop a novel recurrence prediction system based on long-term PCa prognosis instead of interobserver-based histology. We relied on the tissue microarray (TMA) framework of the Canadian Prostate Cancer Biomarker Network (CPCBN) initiative of the Terry Fox Research Institute; this initiative implemented thoroughly validated techniques to ensure the collection of representative samples of PCa from radical prostatectomy (RP) specimens^15^.

In this study, we developed a calibrated and interpretable algorithm for predicting PCa outcomes in multiple independent cohorts that could eventually be integrated into existing prognostic and predictive nomograms.

## Results

### Survival Modeling

To establish a novel system for predicting recurrence, we initially investigated a multicenter population (CPBCN, n = 1,489) in which the overall BCR probability was 33.1% (n = 493). The median time to BCR events was 26 (interquartile range [IQR], 8–52) months; in contrast, the median follow-up was 109 (76– 141) months in patients without BCR events. The development and first external validation sets (CPBCN cohort) were not statistically different with respect to pathological tumor (pT) stage, pathological nodal (pN) status, and GG (Supplementary Table S1). Among 600 patients in the development set, 225 (37.5%) experienced recurrence during follow-up (median follow-up, 91 [42–123] months); in contrast, among 889 patients in the first external validation set, 268 (30.1%) had BCR (median follow-up, 75 [43–116] months).

Fig. 1 summarizes the study methodology using histology images as data input, the confidence scores for BCR as output, and the binarized recurrence status as the ground truth for model development and evaluation. The Supplementary Materials include cohort descriptions for all datasets included in this study (Supplementary Tables S1-S3).

**Fig. 1:**
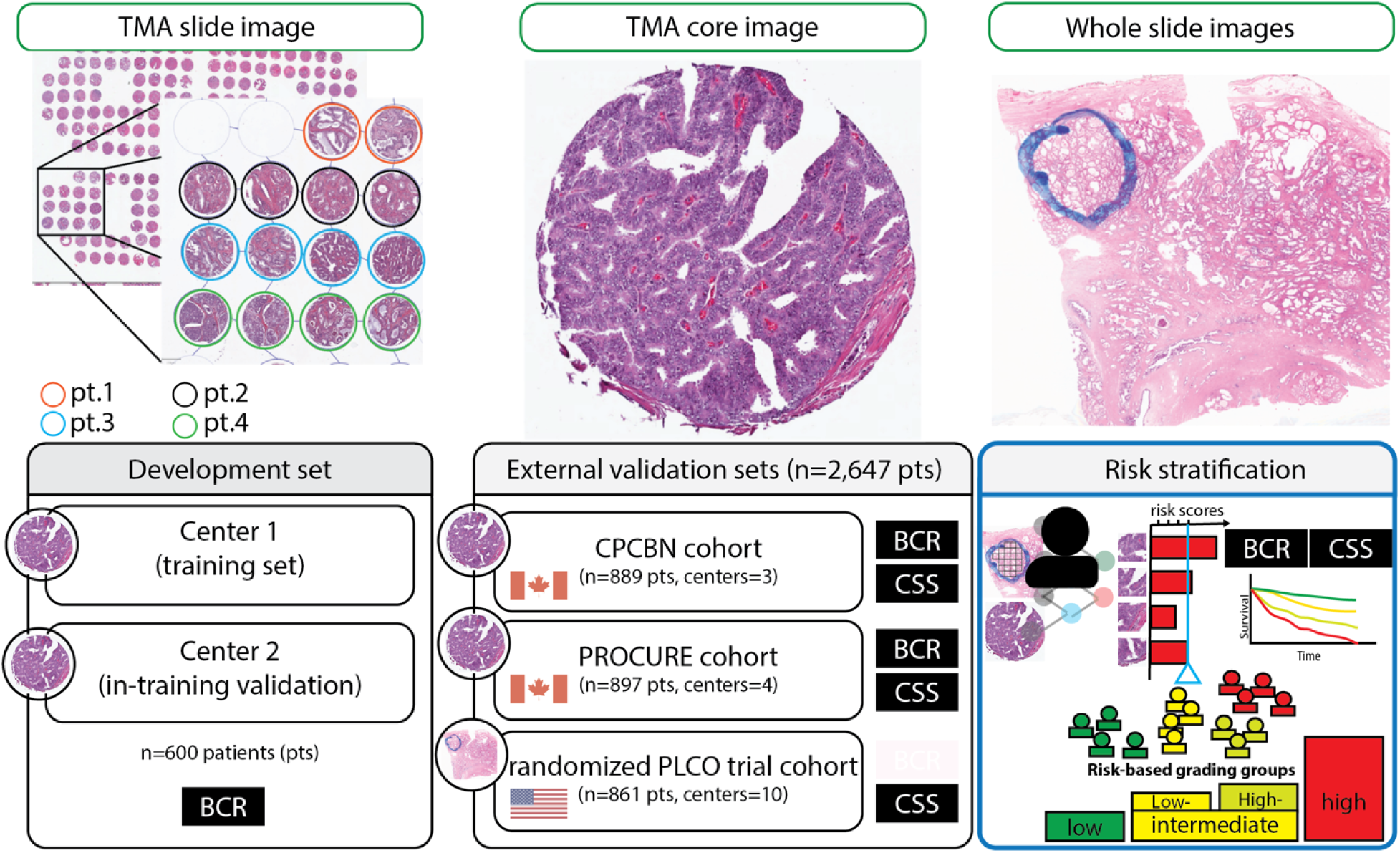
Slides from tissue microarrays (TMAs) with prostates samples from five sites were scanned, and the tissue regions were marked and extracted using QuPath (i.e., TMA slide image). We then tiled each TMA core image into patches labeled by biochemical recurrence (BCR) status to develop our BCR model. We estimated the average BCR scores for each patient and applied survival modeling to introduce our novel risk-based grading for prostate cancer. The development set consisted of 600 patients, whereas the international external validation sets included three radical prostatectomy cohorts (CPCBN, PROCURE, and PLCO). The cohort description for all data sets included in this study can be obtained from Supplementary Tables S1-S3. PLCO: The Prostate, Lung, Colorectal, and Ovarian (PLCO) Cancer Screening Trial. The endpoints we are shown in the black box. CSS: Cancer-specific survival. We emphasize that PC regions were manually demarcated on whole slide images following the instruction given by a senior pathologist.

In the first external validation set, the BCR model demonstrated a c-index of 0.682 ± 0.018 and a generalized concordance probability of 0.927 (95% CI: 0.891–0.952). The AUROC for the BCR model was 0.714 (95% CI: 0.673–0.752). Using a cutoff of 0.5 for the BCR confidence score, the sensitivity was 50.0% and the specificity was 83.2%. The precision and recall of the BCR model at a 0.5 threshold were 56.3% and 50.0%, respectively. The calibration plot demonstrated good correlation between the predicted BCR probability (BCR score) and observed 10-year BCR-free survival rate (Supplementary Fig. 1).

Our novel model revealed a better effect size (hazard ratio) and higher generalized concordance probability than the classical models ResNet^16^, VGG-16^17^, and EfficientNet^18^, which were trained on the same development set for BCR prognosis. EfficientNet and the novel model provided the lowest AIC and BIC. A non-nested partial likelihood ratio test revealed that EfficientNet did not fit better than the novel model. Importantly, our novel BCR model had between 8- and 32-times fewer feature maps in the last convolutional layer for BCR prediction (before being fully connected) and a parameter capacity 125, 54-, or 24-times smaller than the models mentioned above (Supplementary Table S4). We observed no performance benefits from using image patches at 20× or 40× object magnifications, the attention aggregation layer, or the Cox deep convolutional model concept (Supplementary Table S5).

The results of the CHAID analysis are shown in Supplementary Fig. S2. Based on the BCR scores estimated by our model and CHAID, BCR scores ≤5% were considered low risk, BCR scores between 6% and 42% were low intermediate, BCR scores between 43% and 74% were high intermediate, and BCR scores ≥75% were high risk.

### Recurrence-free Survival

One study conducted univariate and multivariable Cox regression analyses on CPCBN and PROCURE cohorts to assess the prognostic value of the novel risk classification system for PCa recurrence (Supplementary Table S6 and S7). The results showed that the BCR score was an independent prognostic factor for recurrence, along with PSA level, tumor stage, GG, and surgical margin status. The novel risk classification system showed a better model fit and superiority over GG (Table 1). No significant multicollinearity between variables was identified (VIF <2), indicating the correlation between variables (GG and the novel risk group) is negligible small.

**Table 1:**
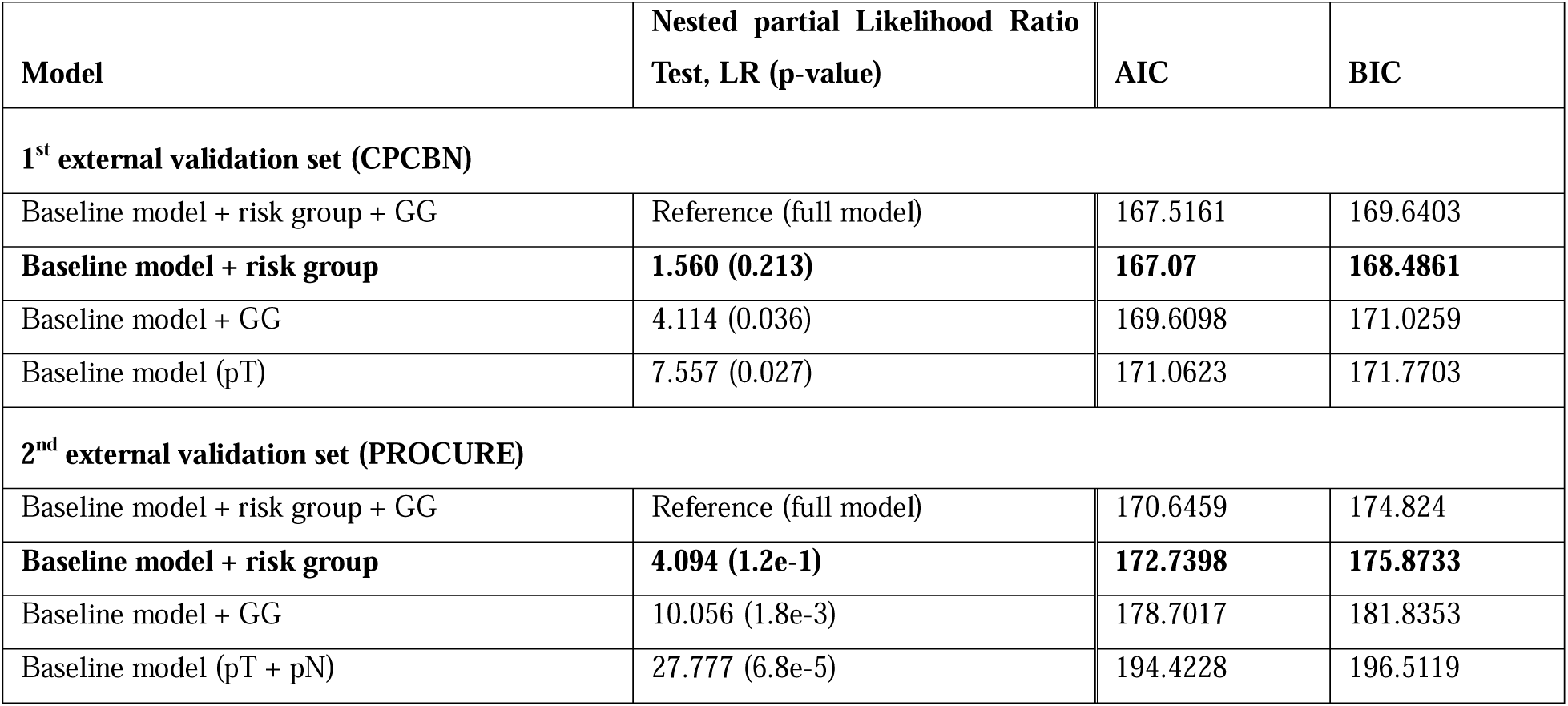
The model reduction and the partial Likelihood Ratio (LR) test revealed that a baseline model with the novel risk groups is statistically comparable to the full model to predict cancer-specific survival. In constrast, GG (Gleason score/ISUP grade groups) was not comparable to the full model. Akaike information criterion (AIC) and Bayesian information criterion (BIC) support this finding as well since the fit of a baseline model with the novel risk groups is better than the fit of a baseline model with GG. pT: pathologic tumor stage; pN: pathologic nodal stage. + pN was excluded due to non-signficance to prognose cancer-specific survival in the CPCBN external validation set. The best performing models are highlighted in bold. Higher AIC and BIC are associated with the worst model fitness. No significant multicollinearity between variables was identified (the Variance Inflation Factors,VIF, were below 2).

The survival rates varied across the novel risk groups in both the cohorts, as shown in and Figures 2A-B (See supplementary Table S8 for 3-, 5-, 10-years BCR-free survival rates). The survival rates for GG are shown in the Supplementary section for comparison (Supplementary Tables S9 and S10 and Figs. S3 and S4). The estimated power for BCR survival analysis in this study was determined to be ≥99% at an alpha level of 5% for each cohort.

**Fig. 2A:**
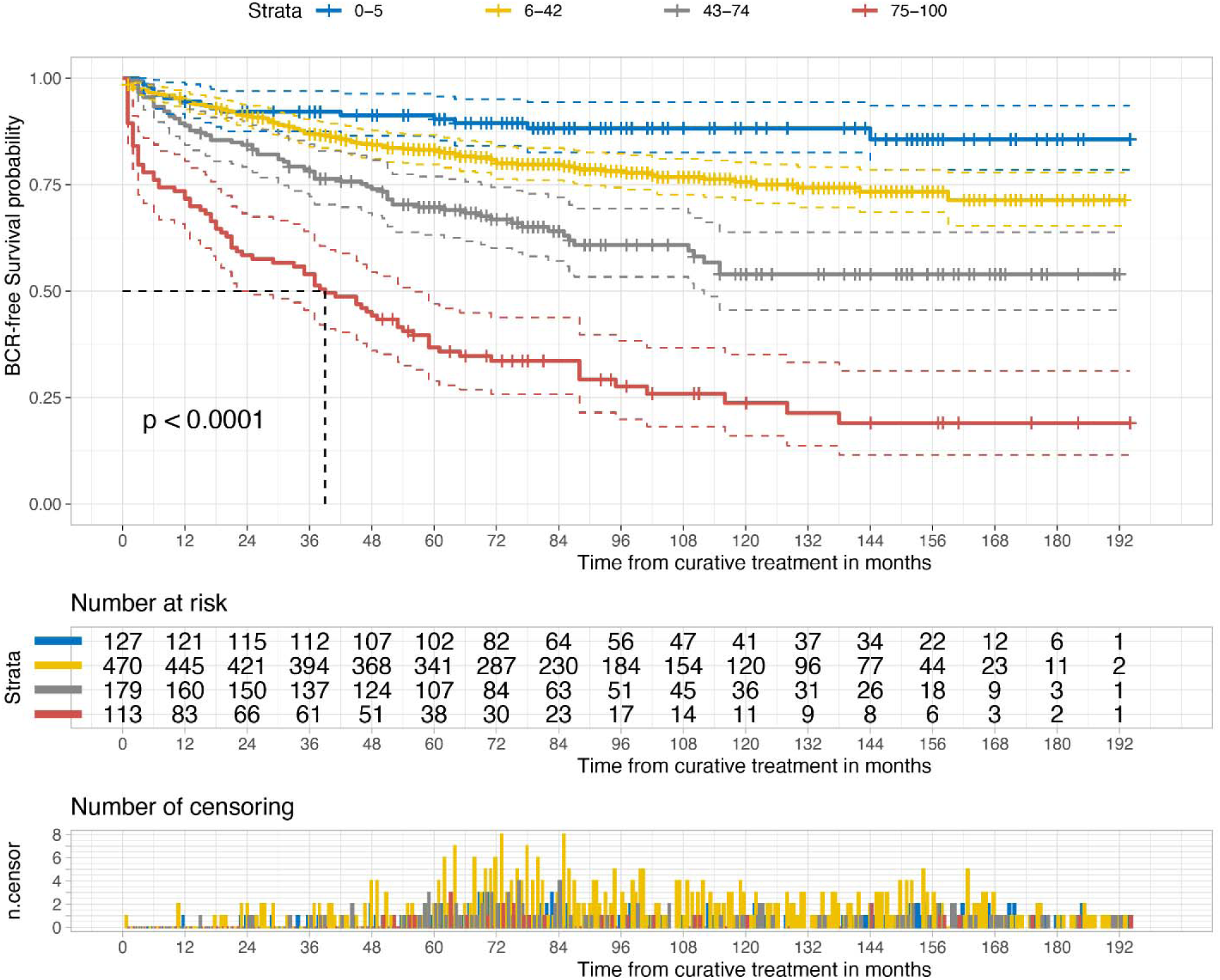
Kaplan-Meier curves of biochemical recurrence (BCR)-free survival according to BCR score risk stratification in the first external Validation set (CPCBN, Canada). P-value was measured using the log-rank test. Blue represents the low-risk group (0-5% BCR score), yellow represents the low-intermediate risk group (6-42%), grey represents the high-intermediate risk group (43-74%), and red represents the high-risk group (75-100%). The dotted lines indicate the median survival. In addition, the number of patients at risk and of censored observations are provided for the follow-up period.

**Fig. 2B:**
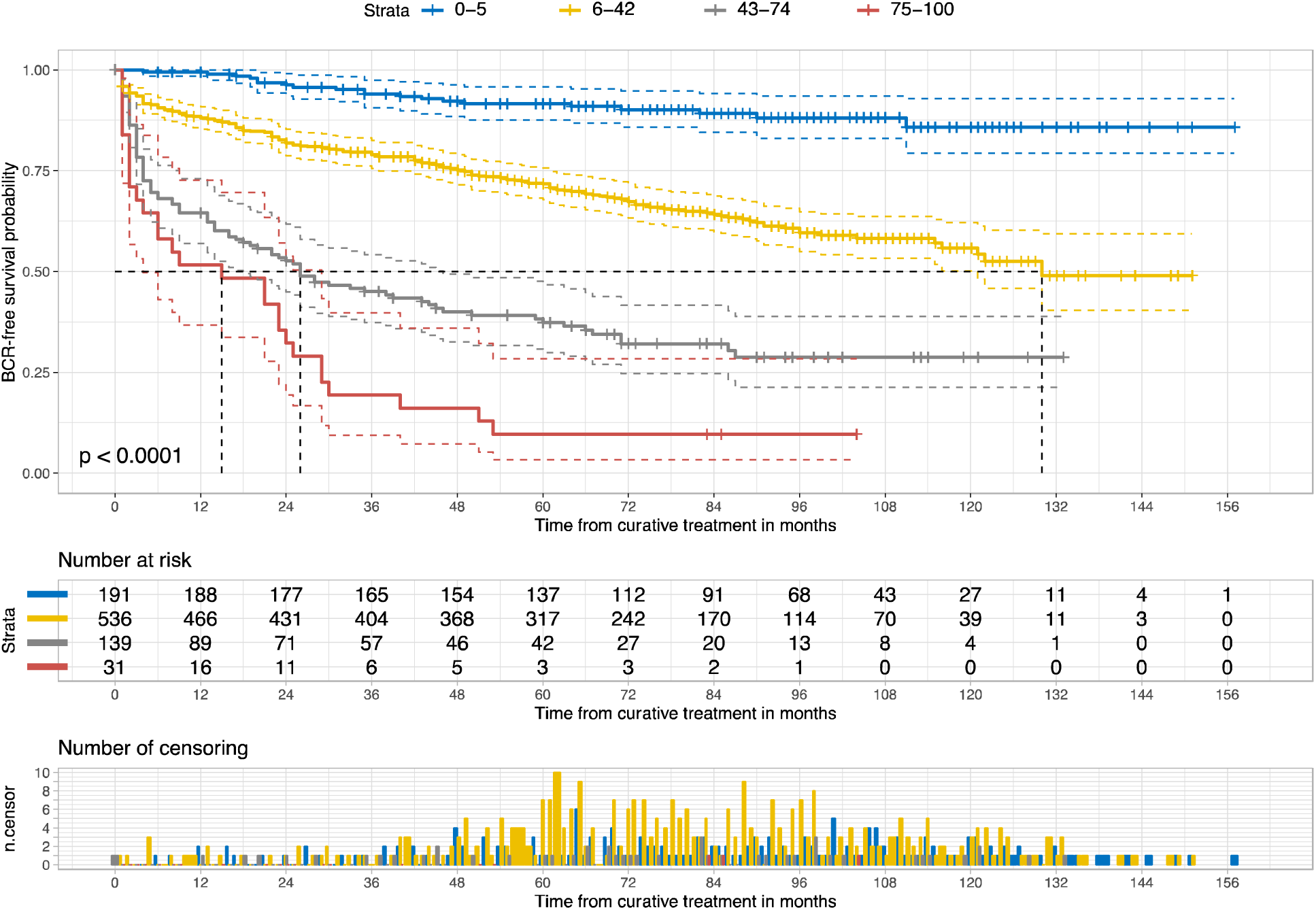
Kaplan-Meier curves of biochemical recurrence (BCR)-free survival according to risk groups in the second external Validation set (PROCURE). Blue represents the low-risk group (0-5% BCR score), yellow represents the low-intermediate risk group (6-42%), grey represents the high-intermediate risk group (43-74%), and red represents the high-risk group (75-100%). The p-value was measured using the log-rank test. The number of patients at risk and of censored observations are provided for the follow-up period.

### Cancer-specific Survival

This study examined cancer-specific survival using a novel risk classification system in three cohorts: the CPCBN, PROCURE, and PLCO cohorts. In the CPCBN cohort, the novel score was a significant prognostic factor for cancer-specific mortality and tumor stage; in contrast, GG was not an independent prognostic factor (Supplementary Table S11). In the PROCURE Quebec Prostate Cancer Biobank (PROCURE cohort), the novel risk score was an independent prognostic factor, along with the nodal stage; in contrast, the tumor stage was insignificant (Supplementary Table S12). Supplementary Table S13 summarizes the results of the Cox regression analyses of the PLCO cohort, further validating the independent prognostic value of the risk score for cancer-specific mortality using whole-slide images.

In the CPCBN and PROCURE cohorts, the multivariate Cox regression model with novel risk groups fit well, similar to the full model. However, the model with GG fits the data poorly (Table 2). In the PLCO cohort, both the GG and risk groups fit poorly compared with the full model, and the difference in the goodness-of-fit between the model with GG and the model with risk groups was insignificant. No significant multicollinearity between variables was identified (VIF <2). The estimated power for BCR survival analysis in this study was determined to be ≥95% at an alpha level of 5% for each cohort. The Fine-Gray competing risk regression analyses further validated the independent prognostic value of our novel risk groups for cancer-specific mortality on external validation sets (Supplementary Tables S14 – S16).

**Table 2:**
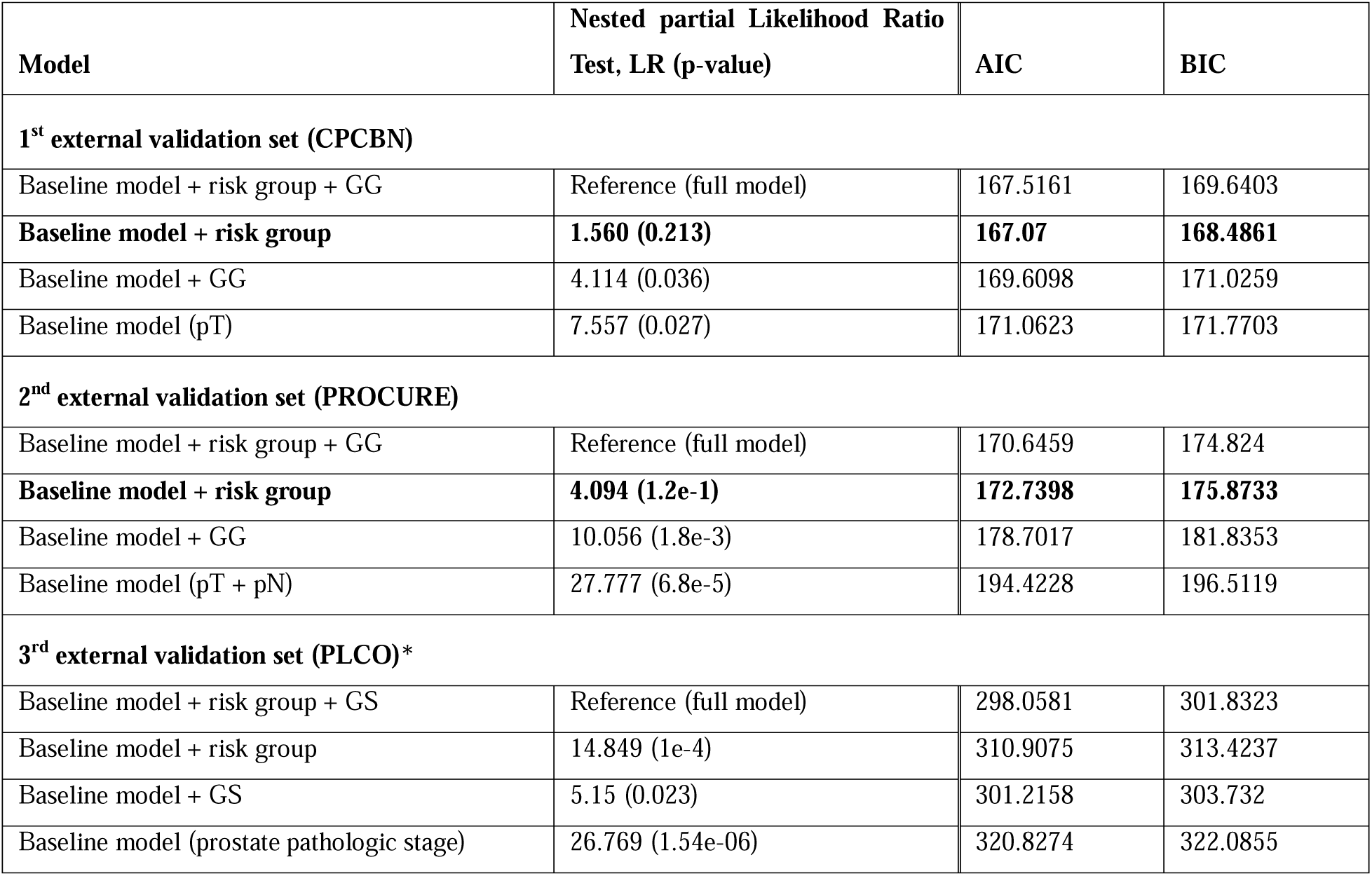
The model reduction and the partial Likelihood Ratio (LR) test revealed that a baseline model with the novel risk groups is statistically comparable to the full model to predict cancer-specific survival. In constrast, GG (Gleason score/ISUP grade groups) was not comparable to the full model. Akaike information criterion (AIC) and Bayesian information criterion (BIC) support this finding as well since the fit of a baseline model with the novel risk groups is better than the fit of a baseline model with GG. pT: pathologic tumor stage; pN: pathologic nodal stage. + pN was excluded due to non-signficance to prognose cancer-specific survival in the CPCBN external validation set. For PLCO external validation set, we used GS provided by the study in stead of GG and prostate pathologic stage (considers T, N and M stages) due to the study history. The best performing models are highlighted in bold. Higher AIC and BIC are associated with the worst model fitness. <u>* since both GS and risk groups were signficanlty inferior than the full model, we</u> <u>applied the non-nested partial likelihood ratio test to compare between GS Cox model and risk group Cox model; our risk</u> <u>group demonstrated non-inferiority to GS, indicating comparable goodness of fit (z = 1.091, p = 0.138).</u> No significant multicollinearity between variables was identified (VIF <2).

The Kaplan-Meier curves for cancer-specific survival according to risk classification in the three external validation sets showed significant differences among the risk groups (Figs. 2C–E). Supplementary Table S17 summarizes cancer-specific survival rates across the three cohorts and shows a distinct separation of survival rates among the risk groups 10 or 15 years after RP. The low-risk group of the novel grading system had no PCa-related deaths in any of the three cohorts; in contrast, the GG in the current grading system included patients who died owing to PCa in two of the three cohorts.

**Fig. 2C:**
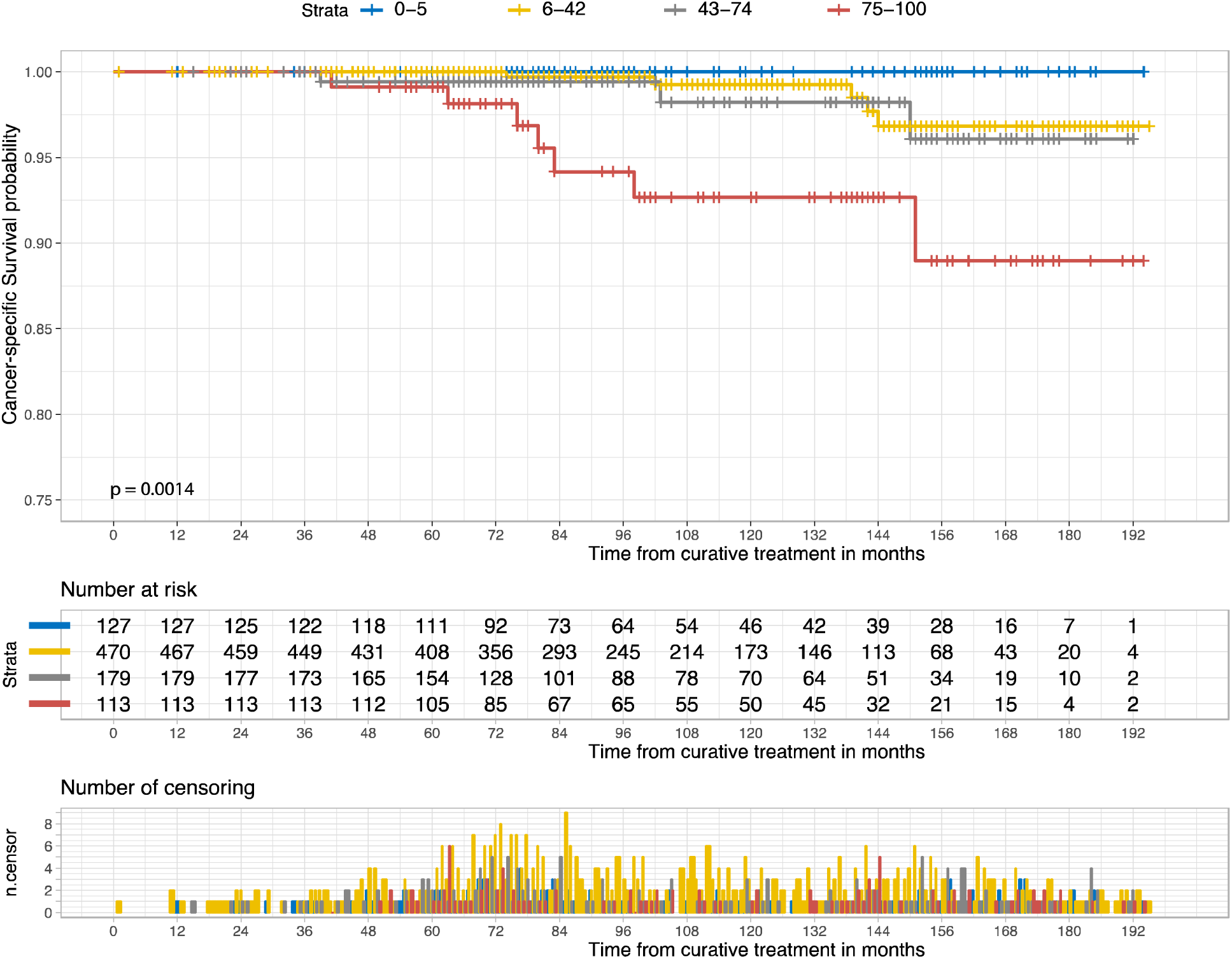
Kaplan-Meier curves of cancer-specific survival according to the risk groups in the first external Validation set (CPCBN, Canada). The P-value was measured using the log-rank test. Blue represents the low-risk group (0-5% BCR score), yellow represents the low-intermediate risk group (6-42%), grey represents the high-intermediate risk group (43- 74%), and red represents the high-risk group (75-100%). The number of patients at risk and of censored observations are provided for the follow-up period.

**Fig. 2D:**
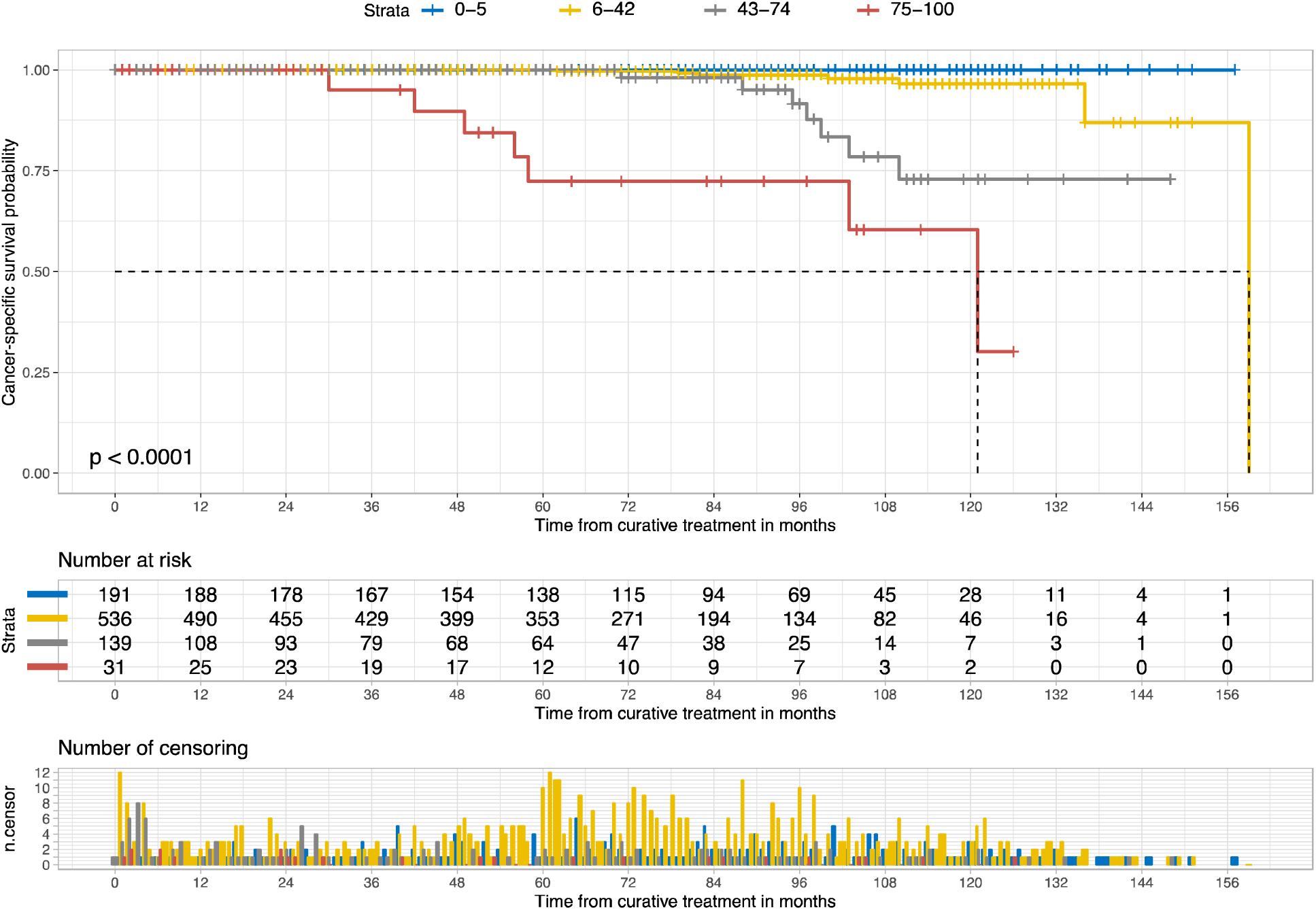
Kaplan-Meier curves of cancer-specific survival according to risk groups in the second external Validation set (PROCURE, Canada). The p-value was measured using the log-rank test. Blue represents the low-risk group (0-5% biochemical recurrence score), yellow represents the low-intermediate risk group (6-42%), grey represents the high-intermediate risk group (43-74%), and red represents the high-risk group (75-100%). The number of patients at risk and of censored observations are provided for the follow-up period.

**Fig. 2E:**
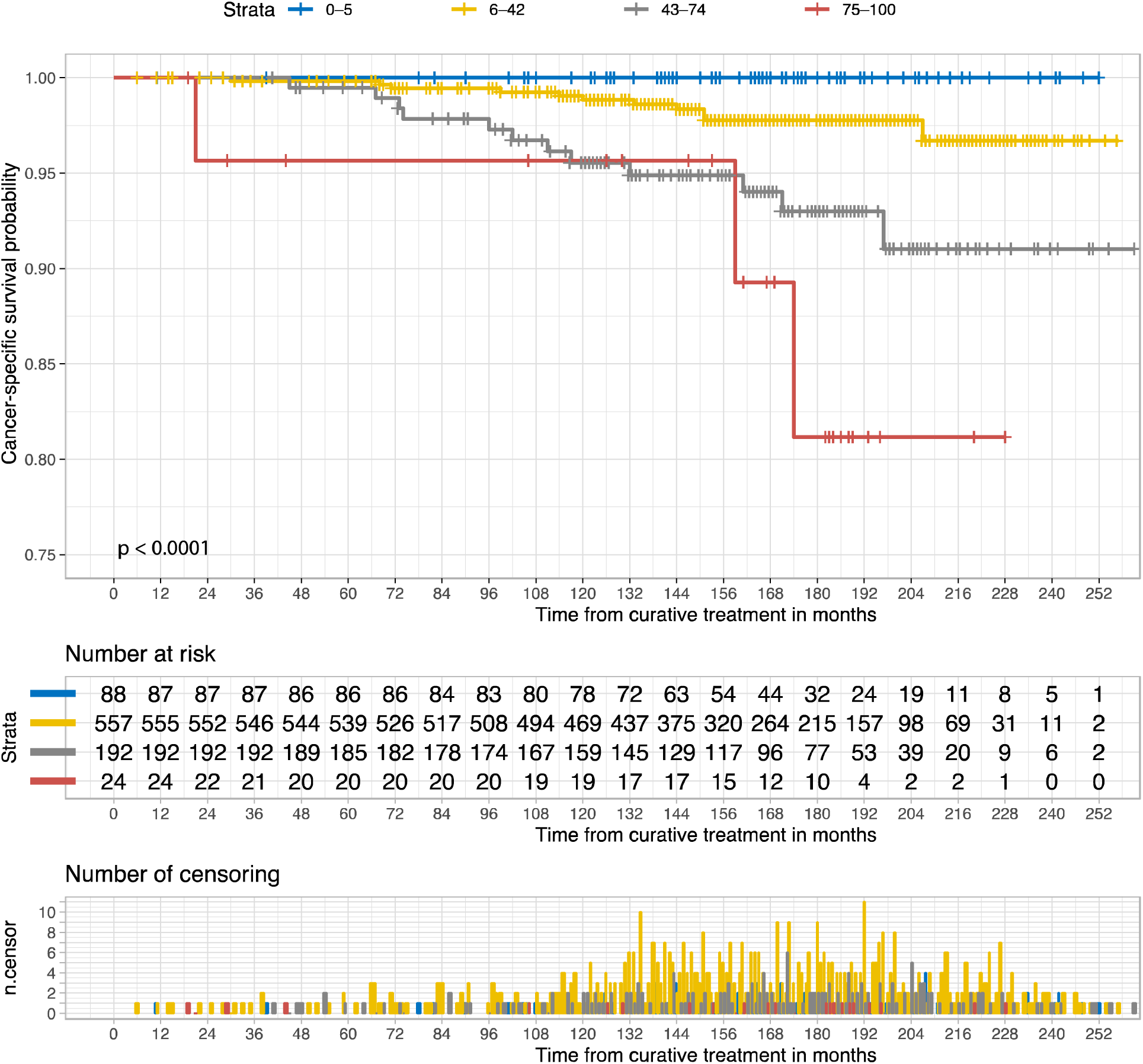
Kaplan-Meier curve of cancer-specific survival according to risk groups in the third external Validation set (PLCO, U.S.). The p-value was measured using the log-rank test. Blue represents the low-risk group (0-5% BCR score), yellow represents the low-intermediate risk group (6-42%), grey represents the high-intermediate risk group (43-74%), and red represents the high-risk group (75-100%). The number of patients at risk and of censored observations are provided for the follow-up period.

PLCO cohort analysis showed that the number of slides per case and its correlation with the risk score did not significantly affect the prognostic value (Supplementary Table S18). Additional information on survival probabilities, Kaplan-Meier curves for the GG, Gleason score groups, and the PCa pathological stage is provided in Supplementary Tables S19–S21 and Supplementary Figures S5–S8 for comparison.

### Interpretability

Table 3 shows the concordance between the five pathologists and novel risk classifications. This table summarizes the synergistic efforts between AI and pathologists in defining a novel grading system for PCa. Despite being completely blinded to the novel risk classification and clinicopathological information, we found a striking alignment between the pathologists and risk classification in sorting image clusters. Despite not relying on pattern proportions like the GG and the absent of significant collinearity between our novel risk group and GG, the image cluster representing the low-risk group included Gleason pattern 3 mostly; in contrast, the high-risk group included Gleason patterns 4 and 5, with Gleason pattern 3 being almost absent. The pathologists found a mixture of Gleason patterns 3 and 4 in the intermediate group, with a trend in favor of Gleason pattern 4 in the high-intermediate group. Fig. 3 exemplary illustrates the histopathological gradient for distortion of glandular architecture as well as the Supplementary section include information on accessing image clusters.

**Figure 3:**
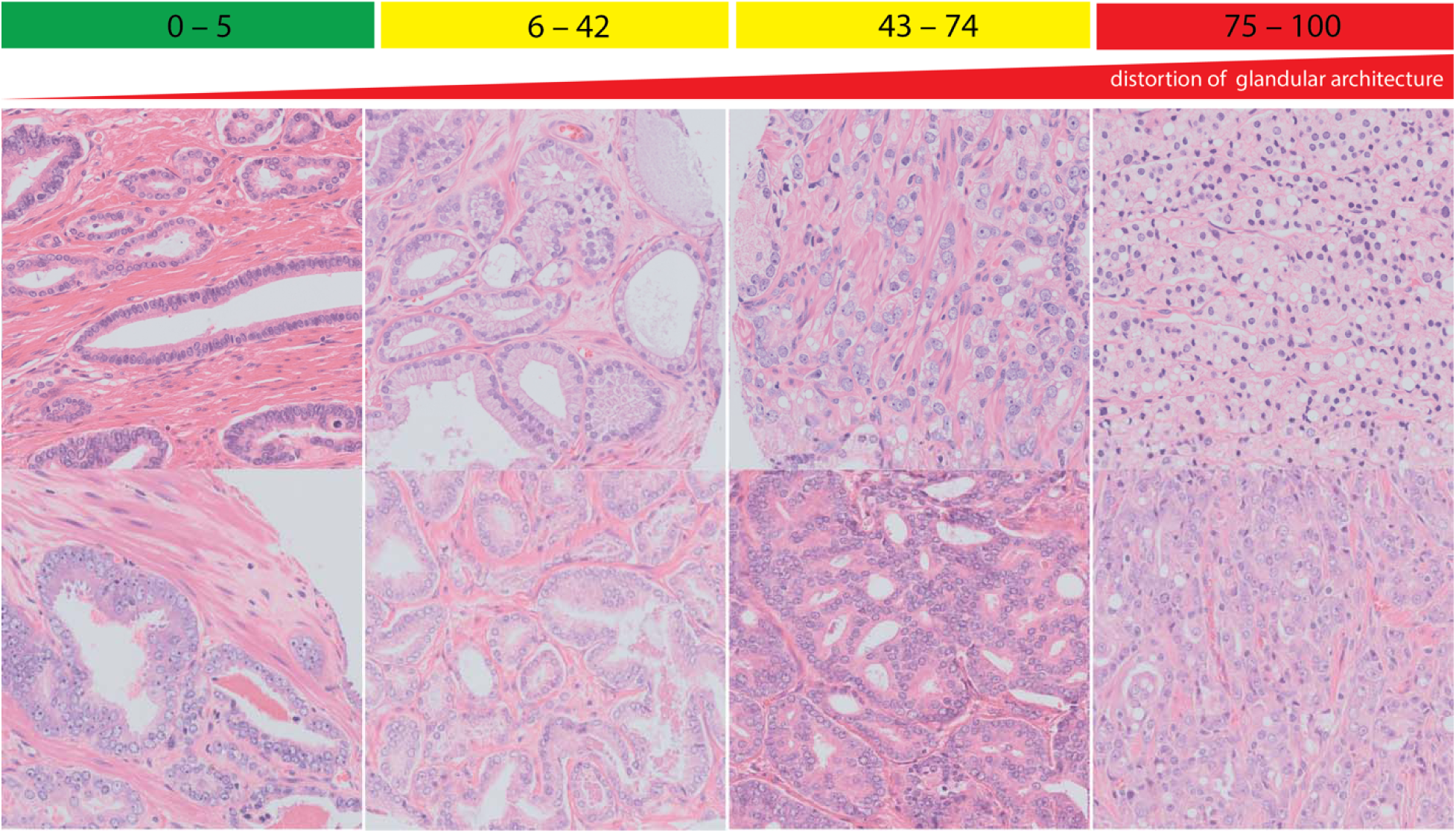
we identified a clear histopathological gradient for distortion of glandular architecture (e.g., disappearance of organized glandular architecture) according to risk groups based on image patches. Example histology images were captured at 10x objective magnification (∼330 × 330 µm). The supplementary section includes the access information to larger image sets representing these risk groups.

**Table 3:** Image cluster assessment by pathologist in accordance with the BCR score risk stratification with decision explanation. Pathologist D weighted the heterogeneity definition to rank Cluster D and C, thereby resulting into the partial agreement.

| Pathologist | Sorting agreement with AI | Explanation |
| --- | --- | --- |
| Pathologist A | Yes | Cluster A: Dominant Gleason pattern 3<br>Cluster B: Mixed Gleason pattern 3 and 4, cannot determine which pattern is dominant.<br>Cluster C: Mixed Gleason pattern 3 and 4, but Gleason pattern 4 is dominant.<br>Cluster D: Few Gleason pattern 3, mostly Gleason pattern 4 and/or 5. |
| Pathologist B | Yes | Cluster A has the most favorable malignancy grade (mostly Gleason pattern 3), whereas Cluster D has the worst differentiation grade (hard to identify Gleason pattern 3, mostly Gleason pattern 4 and 5). Cannot determine a significant difference in pattern distribution in Cluster B and C, but it seems to me that C has more Gleason pattern 4. |
| Pathologist C | Yes | There is a pattern trend in these clusters, likely driven by Gleason pattern 3. For example, Cluster A has mostly Gleason pattern 3 and Cluster D has mostly Gleason pattern 4 and 5. This trend is, however, not clearly visible in cluster B and C (mixed Gleason pattern 3 and 4). |
| Pathologist D | Partially | Cluster A: Dominantly favorable Gleason pattern 3<br>Cluster B: Mixed Gleason patterns 3 and 4<br>Cluster D: Dominantly Gleason pattern 4 and 5 (Mostly Gleason score at least 8)<br>Cluster C: Mostly heterogenous prostate cancer constellation with dominance of Gleason pattern 4. |
| Pathologist E | Yes | Cluster A: Dominant Gleason pattern 3. Rare occurrences of mucinous cribriform, glomeruloid patterns<br>Cluster B: Mixed Gleason pattern 3 and 4, with rare occurrence of single cells (pattern 5). When pattern 4 is present, it is mostly ill-defined, sometimes cribriform.<br>Cluster C: Dominant pattern 4, with rare occurrences of pure pattern 3. Pattern 4 is often cribriform.<br>Cluster D: Dominant patterns 4 and 5. |

## Discussion

In this study, we developed and externally validated a novel grading system for PCa that was superior to the existing grading systems. We demonstrated that AI could be a helpful tool for generating a well-calibrated grading system interpretable by human experts, including risk stratification groups with distinct survival probabilities that enable communication with and between domain experts and between patients and experts to make clinical decisions^7,19,20^. A well-calibrated deep learning model alleviates the usual concerns of overconfidence and enables the interpretation of the model’s prediction as scores^21,22^. Lastly, risk stratification further enables the exploration of common histopathologic patterns by risk scores^7,19,20^. Previous AI efforts have focused on replicating grading systems using supervised learning. Bulten *et al*. reported a deep learning model trained with the semi-automatic region-level annotation technique and slide-level annotations to show a Cohen’s quadratic kappa score (κ_quad_) of 0.918 (95% CI 0.891–0.941)^11^. Similarly, Ström *et al*. developed an ensemble of deep learning models trained with automatically generated region-level annotations from pen marks and slide-level annotations, yielding a linear-weighted kappa score (κ_lin_) of 0.83^23^.

A recent study proposed a weakly supervised deep learning model that leveraged only the global Gleason score of whole-slide images during training to grade patch-pixel-level patterns and perform slide-level scoring accurately^24^. The authors reported an average improvement on Cohen’s quadratic kappa score (κ_quad_) of approximately 18% compared to full supervision for the patch-level Gleason grading task^24^. Similarly, another study reported that the use of the AI-assisted method was associated with significant improvements in the concordance of PCa grading and quantification between pathologists: pathologists 1 and 2 had 90.1% agreement using the AI-assisted method vs. 84.0% agreement using the manual method (p < 0.001)^25^.

Despite these results being promising, the current grading system still suffers from reader dependency, and any AI-based solution developed to improve the interrater agreement for tumor grading will apply to a closed network of human readers with associated social and cognitive biases. To address these integral notions of AI design, our grading system was calibrated with different risk groups independent of human readers. Our approach also overcomes the challenges of interpreting an AI-designed grading system as human readers can identify pattern trends in our grading system. Finally, our novel grading system accurately facilitated PCa grading at the clinically relevant case level using a limited number of representative PCa tissues (three to four small regions representing the index PCa on an RP specimen) or a fully representative slide from an RP specimen.

Previous studies have explored the potential of digital biomarkers or AI-based Gleason grading systems for survival prediction and prognosis in PCa. For instance, a most recent nested case-control study developed a prognostic biomarker for BCR using ResNet-50D^26^ and a TMA cohort, and the time to recurrence was utilized to label the histology images^27^. Wulczyn *et al*. proposed an AI-based Gleason grading system for PCa-specific mortality based on Inception^12^-derived architecture^28^. Yamamoto *et al*. utilized deep autoencoders^29^ to extract key features that were then fed into a second machine learning model (regression and support vector machine^30^) to predict the BCR status for PCa at fixed follow-up time points (Year 1 and 5)^31^. Other studies also utilized multimodal data (molecular feature and histology) for prognosis in different cancers^32,33^. Overall, these studies set the ground for further survival analyses using AI; however, they were limited by the post hoc explanation of their black box models that is not necessarily reflective of interpretable, clinically relevant well-validated algorithms^34–36^.

One of the most important aspects to consider when developing tools for clinical decision-making is practicality and clinical utility. Our novel model was calibrated to predict 10-year BCR-free survival probability and facilitate model interpretation. It should also be noted that the standard prognostic factors for PCa are all obtained during diagnosis or treatment without accounting for any time information.

Accordingly, we integrated this important aspect into our novel prediction system and selected model architectures for comparison based on recent surveys for medical imaging^37,38^ and the PANDA Challenge^39^ for PCa. Similarly, because c-index and ROC curves are not ideal for comparing prognostic models, we utilized the partial LR test, AIC, and BIC to identify which model configuration fits better and provides a superior prognostic performance^40^. The novel prediction system presented in this study does not rely on Cox models to calculate risk scores and determine risk groups. In this study, Cox models were used only to evaluate the accuracy and clinical utility of the grading system.

This study applied the Gleason grading system for nomology and ontology to describe the histopathological contents of each group as it is widely accepted as a communication terminology for histopathological changes in PCa among domain experts (including urologists, pathologists, and oncologists), despite their interrater limitations. Although there was some unsurprising overlap between our risk scores and the GG, the risk groups provided significantly different interpretations of the GG patterns. Furthermore, our analysis revealed no significant evidence of multicollinearity among various parameters, including Gleason grade (GG) and the risk groups. This suggests that the variables we considered in our study are independent and not significantly correlated with each other.

Although our results are robust, and our novel grading system does not rely on GG nor pattern proportions, whether it can overcome sampling errors, tissue fragmentation, degradation, or artifacts caused by prostate biopsy and/or poor RP tissue quality is unknown. We did not evaluate our grading system on the biopsy materials for survival modeling as a sampling effect (evident from the increase in PCa on RP) and the effects of time or intermediate events (such as cancer progression) until treatment (such as RP) were difficult to control in the experimental setting. In contrast, these effects were easier to control with RP specimens, and it was previously demonstrated that TMA, corresponding biopsy samples and RP specimens were comparable to GG^15,41^. The selection strategy for whole-slide images (WSIs) or tissue microarray (TMA) sampling in the current cohorts was determined exclusively by the study organizers before the initiation of the current study. Thus, our strategy mitigated the observer bias by ensuring that data collectors were not involved with data analysis process of the current study. Although we did not have control over the WSI or TMA sampling and case selection process for the current study, our power analyses indicate that the sample size we have is adequate to execute our study. Moreover, the TMA cohorts were primarily designed for biomarker validation, specifically to assess the effectiveness of biomarkers in predicting or prognosing survival outcomes. The selection of TMA samples accordingly followed predetermined criteria set by the study organizers to ensure accurate representation and robust validation while mitigating the selection bias ^15,41^. To mitigate potential bias from interobserver variability in labeling histopathological image clusters, we requested explanations from pathologists to better understand the factors influencing their decisions. This approach aimed to improve transparency and provide insights into the potential sources of bias in the interpretation of histopathological images. Finally, our AI-based grading system was not developed to detect PCa; therefore, additional models to detect PCa are required for a fully automated grading system.

This study introduced and validated a novel grading system resulting from the synergy between AI and domain knowledge. Future research should focus on identifying the application boundaries of our novel grading system in a real-world setting, including its possible integration into existing nomograms used to predict prognosis and treatment response.

## Data Availability

Due to data transfer agreements and data privacy issues, data cannot be made openly available.

## Code Availability

An abstract version of the codes can be obtained from https://github.com/oeminaga/AI_PCA_GRADE.

## Acknowledgements

The Canadian Prostate Cancer Biomarker Network (CPCBN) acknowledges contributions to its biobank from several Institutions across Canada: Centre hospitalier de l’Université de Montreal (CHUM), Centre hospitalier universitaire de Quebec (CHUQ), McGill University Health Centre (MUHC), University Health Network (UHN), and University of British Columbia/Vancouver Coastal Health Authority. D.T. receives salary support from the FRQS (Clinical Research Scholar, Junior 2). The CRCHUM and CRCHUQc-UL receive support from the FRQS. The authors thank Mrs. Véronique Barrès, Mrs. Gabriela Fragoso, and Mrs. Liliane Meunier of the molecular pathology core facility of the Centre de Recherche du Centre hospitalier de l’Université de Montréal for performing the sections, immunohistochemistry, and slide scanning and the facility core for image analysis with the Visiopharm image software. Access to Dr. Féryel Azzi’s expertise is possible thanks to the TransMedTech Institute and its primary funding partner, the Canada First Research Excellence Fund. We acknowledge the contribution of PLCO study trial to this study providing histology images and corresponding clinical data. Finally, external validation TMAs for this research project were obtained from the PROCURE Biobank. This biobank is the result of a collaboration between the Centre hospitalier de l’Université de Montréal (CHUM), the CIUSSS de l’Estrie-CHUS, the CHU de Québec-Université Laval and the Research Institute of the McGill University Health Center, with funds from PROCURE and its partners. We thank the organization of the PLCO study for sharing the whole-slide images and the corresponding clinical information.

## Online Methods

### Data

#### Cohorts

In this study, we adopted a study design that focused on the analysis of independent retrospective cohorts. The development cohort included 600 RP cases from two institutions in the CPCBN framework^15,41^. The first external validation set, the CPCBN cohort, included 889 RP cases from three different institutions within the CPCBN framework, anonymized to minimize bias and excluding the institutions used in the development set to avoid potential label leakage. The second cohort included 16 digital TMA scans of 897 patients from the PROCURE cohort^42,43^. Lastly, the 1,502 H&E-stained whole-slide images from 861 RP cases in the Prostate, Lung, Colorectal, and Ovarian (PLCO) Cancer Screening Trial (NCT00339495; PLCO cohort) were used^44,45^. Only cores or representative slides from the RP index lesion were used to develop and validate the malignancy grading system for PCa. The Supplementary Methods details TMA construction and histological images of these cohorts as well as their exclusion and inclusion criteria.

#### Clinicopathological Information

Histological images of PCa, clinicopathological information, and longitudinal follow-up data were available for all cases. Clinicopathological data included age at diagnosis, preoperative prostate-specific antigen (PSA) measurements, RP TNM classification, and RP GG for all patients at the RP and TMA core sample levels. Tumor staging was based on the 2002 TNM classification^46^ and grading according to the 2016 WHO/ISUP consensus^47^. All data were available from the corresponding framework and study trial. The clinicopathological information was obtained through a meticulous chart review process, involving the extraction of data and the data quality control from the electronic health records (EHR) of each participating hospital.

#### Follow-up and Endpoints

Most patients were regularly followed after RP to identify BCR, defined as two consecutive increases in serum PSA levels above 0.2 ng/mL, PSA persistence (failure to fall below 0.1 ng/mL), initiation of salvage or adjuvant treatment, and cancer-specific death. BCR status (non-BCR vs. BCR) and cancer-specific death status were documented during the follow-up period. Non-BCR cases or cancer survivors with incomplete follow-up duration were censored at the date of last follow-up for survival analyses.

### Model Development

The development cohort was further divided into training and in-training validation sets, with the largest single-institution cohort used as the training set. Gleason patterns were utilized to ensure consistent histological appearance in circular cores with a diameter of approximately 0.6 mm. Gleason patterns 3 + 3 and 4 + 4 were specifically used to evaluate homogeneous cores to ensure consistency in the histological appearance. These patterns were selected to determine the minimum and maximum ranges of tissue distortion within the circular cores. In contrast, cores with Gleason pattern 4 + 3 were considered to represent heterogeneous cores, indicating an intermediate stage of tissue distortion. The selection of Gleason pattern 3 cores was limited to cases without recurrence during follow-up to ensure a clean pattern. Images including Gleason pattern 5 were intentionally excluded from the training set. By removing pattern 5 and 3+4 from the training set, we aimed to encourage the model to learn and rely on other distinguishing features that are indicative of different malignancy patterns other than the Gleason pattern system (quasi zero-shot learning). As a result, the model development process accounted for tissue appearance and distortion variations independent of the current Gleason grading system.

The study employed neural architecture search using PlexusNET and grid search to find the best architecture model for BCR prediction^48^. ADAM optimization algorithm and cross-entropy loss function were used to train the models^49^. The optimal architecture was selected based on a 3-fold cross-validation performance. The resulting model was trained on the entire training set with early stopping and triangular cyclical learning rates applied to mitigate overfitting. Model performance was evaluated at the case level using confidence scores and metrics such as AUROC and c-index^50,51^. Tile-level predictions were aggregated to determine core- or slide-level predictions, and case-level predictions were estimated by averaging core- or slide-level predictions. The Supplementary Methods section provides more details about model development.

Additional analyses were conducted using other neural network architectures and techniques, as described in the Supplementary Methods. The performance benefits of using different magnifications and survival modeling approaches were assessed. The risk classification model for BCR was constructed using the chi-square automatic interaction detector (CHAID) algorithm^52^, with probabilities cutoffs identified on the development set and validated on external validation sets.

### Model Evaluation

In the development and external validation cohorts, confidence scores for BCR (BCR scores) were generated for all cases. Prognostic classification and accuracy were measured using AUROC, Harrell’s C-index, and generalized concordance probability. The goodness-of-fit was assessed using Akaike information criterion (AIC) and Bayesian information criterion (BIC)^53–55^.

Calibration plots were created for external validation of the BCR model to evaluate its interpretability. Harrell’s “resampling model calibration” algorithm was applied to assess model calibration^56,57^. BCR predictions were compared to corresponding Kaplan-Meier survival estimates within 10 years.

Univariate and multivariate weighted Cox regression analyses were conducted on external validation cohorts using Schemper *et al*.’s method to provide unbiased hazard ratio estimates, even in cases of non-proportional hazards^58^. Parameters included age at diagnosis, surgical margin status, preoperative serum levels of PSA, pT stage, pN stage, GG, and BCR confidence scores. Parameters significant in the univariate analysis were included in the multivariate Cox regression analysis to identify independent prognostic factors for BCR.

Cox regression models were used for cancer-specific survival to examine the prognostic value of the novel score/grading system, including GG, tumor stage, and the novel score/grading system. In addition to that, we performed the Fine-Gray competing risk regression analyses for cancer-specific mortality, while considering other competing causes of death reported in the death certificates. Kaplan-Meier survival estimates were used to approximate the BCR and cancer-specific survival probabilities for GG and the novel risk classification.

Nested partial likelihood ratio tests were conducted to compare different Cox regression model configurations (only categorical variables) and determine the best model for prognosis^59^. The best-performing grading system (novel grading vs. GG) was chosen based on the lowest changes in partial likelihood ratio and p-values. The AIC and BIC values were compared among the Cox regression models, with the best model having the lowest values. Pearson correlation coefficient was calculated to assess the correlation between the risk score and slide number.

The variance inflation factor (VIF) was utilized to assess the multicollinearity level between the GG, novel grading, and tumor stage. Here, we built two logistic regression models for 10-year BCR and cancer-specific death prediction. VIF below 2 indicates a negligible multicollinearity between these prediction variables.

To ensure the robustness, reliability, and adequate sample size of our study, we conducted a power calculation for Cox proportional hazards regression. Specifically, we evaluated the statistical power of our analysis considering GG and risk score groups to prognose BCR or cancer-specific mortality using powerSurvEpi^60^.

### Human interpretability

The first external validation set (CPCBN) images were clustered according to a risk classification model. Five experienced genitourinary pathologists with over 10 years of expertise were asked to review and sort randomly labeled image clusters based on tumor differentiation. Furthermore, these senior pathologists had to explain their decision in sorting the image clusters while no specific instruction on how to explain their decision was given. Pathologists were blinded to the corresponding clinicopathological and follow-up information to mitigate the recall bias and survivorship bias. Each pathologist was individually approached via email to perform the assigned task while the image clusters were randomly sorted before sharing them with each pathologist; no communication between pathologists specific to this task was permitted to avoid the confirmation bias. Time limitation was not set to execute the task. To assess the inter-rater agreement between a pathologist and our novel risk groups, we utilized a percent agreement based on the proportion of correctly labeled risk groups out of the total number of risk groups under the assumption that the probability for a random agreement in sorting the entire clustered images between a single pathologist and the novel risk classification model is <5% and therefore negligible.

### Software

Model development and analyses were performed with Albumentations^61^, Keras 2.6^62^, TensorFlow 2.6^63^, Python™ 3.8, SPSS® 23 and the R statistical package system (R Foundation for Statistical Computing, Vienna, Austria).

## Notes

**Competing Interests** All Authors declare no Competing Financial or Non-Financial Interests.

### Competing Interest Statement

The authors have declared no competing interest.

### Summary of Updates

Updated the affiliation of Okyaz Eminaga to AI Vobis

