## Supplementary material for "Artificial Intelligence Helps to Predict Recurrence and Mortality for Prostate Cancer Using Histology Images"

### Cohort description

#### The image cohort of the Canadian Prostate Cancer Biomarker Network's Framework

Five different biobanks affiliated with Canadian academic health care centers contributed radical prostatectomy (RP) specimens from 1990 and 2011 to the Canadian Prostate Cancer Biomarker Network (**CPCBN**): Centre hospitalier de l'Université de Montréal (CHUM), Center de recherche du CHU de Québec-Université Laval (CHU de Québec-UL), McGill University Health Centre (MUHC), Vancouver Prostate Centre (VPC), and the University Health Network (UHN). All subjects signed informed consent to contribute to one of the participating biobanks, and each institution's ethical board approved this study, and institutional authorization was obtained.[1,2] RP as primary treatment was performed within six months after the primary diagnostic biopsy. The CPCBN biorepository is complemented by data on diagnosis, treatment, and long-term clinical outcomes. Tumor and lymph node staging was performed according to the 2002 TNM classification by the AJCC/UICC (American Joint Committee on Cancer/International Union Against Cancer).[3] Tumor grading was evaluated according to the 2016 WHO/ISUP consensus.[4]

TMAs were constructed and designed from the RP specimens from 1,489 patients using a comprehensive protocol with rigorous tissue quality assessment, including immunohistochemistry staining, independent double central pathology review by genitourinary expert pathologists as described previously, and routine histology slides superimposed against immunohistochemistry targeting basal cells.[1,2] Each TMA block contained three to four cores from the RP index lesion and one to two cores from adjacent benign tissues within the same individual. Our study considered only cores from the RP index lesion to develop a malignancy grading system for prostate cancer (PCa).

#### The image cohort of Quebec Prostate Cancer Biobank (PROCURE)

A total of 16 digital TMA scans representing 897 cases from the prospectively maintained **PROCURE** Quebec Prostate Cancer Biobank were utilized for external validation.[5,6] The biobank includes patients with localized PCa, who were treated with RP between November 2006 and April 2013 in one of the following University hospitals: Laval University, McGill University, Université de Montréal, and Université de Sherbrooke (**Supplementary Table S2**). Our cases were selected to curate a representative cohort with a minimum median follow-up of 5 years and sufficient image quality, i.e., no tissue degradation/fragmentation or tissue damages and ≥50% tumor burden per TMA spot. All TMA slides of the PROCURE cohort were stained with H&E and digitally scanned at 20x objective magnification using a Leica Biosystems device (Wetzlar, Germany) and stored in SVS format. We then utilized QuPath[7] (Dublin, Ireland) to extract core images (1450 × 1450 pixels or 717.75 × 717.75 µm where 0.4950 µm corresponds to one pixel) as described above. Histology images representing PCa and corresponding clinicopathological information and longitudinal follow-up data were available for all cases. Clinicopathological data included age at diagnosis, preoperative prostate-specific antigen (PSA) measurement (ng/mL), RP TNM classification, and RP GGs

##### The image cohort of Prostate, Lung, Colorectal, and Ovarian (PLCO) Cancer Screening Trial

A third external validation set consisted of 1,502 H&E-stained whole slide images from 861 RP cases enrolled in the Prostate, Lung, Colorectal, and Ovarian (PLCO) Cancer Screening Trial (NCT00339495).[8,9] This study used a linkage with the National Death Index to extend mortality follow-up to a maximum of 19 years after randomization.[10] The slides of the RP **PLCO cohort** were obtained from 10 US centers and digitally scanned at 40× objective magnification (one pixel corresponds to 0.2532 µm) using a Leica Biosystems device (Wetzlar, Germany) and stored in SVS format. The cancer lesions were marked on the slides by PLCO pathologists, and a single PCa researcher of our team digitally verified and delineated these lesions using Aperio Image Scope (Leica Biosystems, Wetzlar, Germany). In addition, clinicopathological information was available at the case level.

Histology images representing PCa and corresponding clinicopathological information and longitudinal follow-up data were available for all cases. Clinicopathological data included age at diagnosis, preoperative prostate-specific antigen (PSA) measurement (ng/mL), RP TNM classification, and RP grade groups (GGs) for all PCa patients at RP and at TMA core sample level if applicable.

#### Image extraction from tissue microarray (TMA) slide images

All TMA slides were stained with hematoxylin and eosin (H&E) and digitally scanned at 40× objective magnification using an Olympus scanner device (Tokyo, Japan) and stored in TIFF format. We then utilized QuPath[7] (Dublin, Ireland) to extract core images (4096 × 4096 pixels or 659.58 × 659.58 µm where 0.1613 µm corresponds to one pixel). We followed the instructions provided by Andrew Janowczyk to extract the core (spot) images from the whole slide TMA images (http://www.andrewjanowczyk.com/de-array-a-tissue-microarray-tma-using-qupath-and-python/; accessed 11/2022). Each core was categorized as malignant or benign.

#### Image preprocessing

To avoid unnecessary computational costs of large images, the core images or the marked prostate cancer area on whole slide images were tiled into small patches at 10× objective magnification level (512 × 512 pixels or ~330 × 330 µm) with a 50% overlap rate (i.e., 50% of the information content of two patches were similar). We considered the standard ISO RGB (red, green, and blue) color space with three color channels[11]. Instead of stain color normalization, we applied color space augmentation (i.e., manipulation of saturation, contrast, and brightness) to reduce the effect of color space on the model inference.

#### Supplementary table S1

*Table S1: Cohort characteristics of the study population (CPCBN, Canada). The distribution of age at diagnosis, preoperative PSA level as well as pathological tumor stage and lymph node status did not significantly differ between development and external validation sets (p>0.05). However, we observed a significant association of the distribution of biochemical recurrence status (p=0.00310) and grade group (GG) at radical prostatectomy (RP) (P=0.01963) with the definition of the data set. We emphasize that our model was trained only on 319 cases with grade group I, III and IV, which represent two homogenous grade groups and one heterogenous grade group, as explained in the "Online Methods" section, and which was evaluated on the whole development set and external validation set. IQR: interquartile range; PSA: prostate-specific antigen; CI: confidence interval.*

| | Overall | Development set | External validation set |
|---|---|---|---|
| Patients, n (%) | 1,489 (100) | 600 (40.3) | 889 (59.7) |
| Age at diagnosis, years, median (IQR) | 62 (57–66) | 61 (57–65) | 62 (57–67) |
| Preoperative PSA level, ng/mL, mean (95% CI) | 8.76 (8.32–9.19) | 8.76 (8.19–9.33) | 8.75 (8.13–9.38) |
| Grade groups at RP, n (%) | | | |
| GG1 | 479 (32.2) | 209 (34.8) | 270 (30.4) |
| GG2 | 595 (40.0) | 250 (41.7) | 345 (38.8) |
| GG3 | 224 (15.0) | 77 (12.8) | 147 (16.5) |
| GG4 | 112 (7.5) | 33 (5.5) | 79 (8.9) |
| GG5 | 79 (5.3) | 31 (5.2) | 48 (5.4) |
| Tumor stage, n (%) | | | |
| pT2 | 946 (63.5) | 395 (65.8) | 551 (62.0) |
| pT3a | 382 (25.7) | 152 (25.4) | 230 (25.8) |
| pT3b | 139 (9.3) | 53 (8.8) | 86 (9.7) |
| pT4 | 22 (1.5) | 0 (0.0) | 22 (2.5) |
| Lymph node status, n (%) | | | |
| pN0/X | 1,443 (96.9) | 586 (97.7) | 857 (96.4) |
| pN1 | 46 (3.1) | 14 (3.2) | 32 (3.6) |
| Biochemical recurrence status, n (%) | | | |
| No | 996 (66.9) | 375 (62.5) | 621 (69.9) |
| Yes | 493 (33.1) | 225 (37.5) | 268 (30.1) |
| Prostate Cancer-specific death status, n (%) | | | |
| No | 1,450 (97.4) | 576 (96) | 874 (98.3) |
| Yes | 39(2.6) | 24 (4) | 15 (1.7) |
| Median core number per patient, n (range) | 3 (1–8) | 3 (1–8) | 3 (1 –8) |
| Total cores (images), n (%) | 4,479 (100) | 1,699 (37.9) | 2,780 (62.1) |
| Total patches, n (%) | 40,311 (100) | 15,291 (37.9) | 25,020 (62.1) |
| Number of centers, n (%) | 5 (100) | 2 (40%) | 3 (60%) |

### Supplementary table S2

*Table S2: Cohort characteristics of the study population (PROCURE, Canada). IQR: interquartile range; CI: Confidence interval; CRPC: castration-resistant prostate cancer; RP: Radical prostatectomy.*

| | Overall |
|---|---|
| Patients, n (%) | 897 |
| Age at diagnosis, years, median (IQR) | 62 (58–67) |
| Preoperative PSA level, ng/mL, mean (95% CI) | 7.8 (7.28–8.32) |
| Grade groups at RP, n (%) | |
| GG1 | 143 (15.9) |
| GG2 | 455 (50.7) |
| GG3 | 204 (22.7) |
| GG4 | 27 (3.1) |
| GG5 | 68 (7.6) |
| Tumor stage, n (%) | |
| pT2 | 535 (59.6) |
| pT3a | 252 (28.1) |
| pT3b | 110 (12.3) |
| Lymph node status, n (%) | |
| pN0/X | 858 (95.7) |
| pN1 | 39 (4.3) |
| Biochemical recurrence status, n (%) | |
| No | 568 (63.3) |
| Yes | 329 (36.7) |
| Time to biochemical recurrence in months, median (IQR) | 21 (4–46) |
| Prostate Cancer-specific death status, n (%) | |
| No | 876 (97.7) |
| Yes | 21 (2.3) |
| Time to cancer-specific death in months, median (IQR) | 95 (62–103) |
| Median core number per patient, n (range) | 3 (1-7) |
| Total cores (images), n | 2,673 |
| Total patches, n (%) | 24,057 |
| CRPC development during follow-up, n (%) | 47 |
| Time to CRPC in months, median (IQR) | 70 (53.5–94.0) |
| Number of centers, n (%) | 4 |

### Supplementary table S3

*Table S3: Cohort characteristics of the third external validation set (PLCO, U.S.). Grade grouping according to Epstein et al. was not feasible to determine due to the impossibility to differentiate between Gleason scores 3+4 and 4+3 from the PLCO dataset. AJCC prostate pathologic stage system combines TNM classification system, Gleason score, preoperative PSA levels. No patients had any evidence for distant metastases nor received neoadjuvant hormone therapy. Patients were prospectively randomized for PSA screening and enrolled in 10 multinational U.S. centers. In all slides, PCa lesions were delineated according to the pathologist's instruction. Slides with PCa were representative of the index lesion for each patient. IQR: interquartile range; PSA: prostate-specific antigen; CI: confidence interval; RP: radical prostatectomy.*

| | Overall |
|---|---|
| Patients, n | 861 |
| Age at diagnosis, years, median (IQR) | 61 (58–68) |
| Preoperative PSA level, ng/mL, mean (95% CI) | 6.45 (6.14–6.76) |
| Pathologic tumor stage, n (%) | |
| pT2 | 670 (77.8) |
| pT3a/b | 191 (22.2) |
| Lymph node status, n (%) | |
| pN0/x | 861 (100.0) |
| Gleason score at RP | |
| Gleason score 6 | 443 (51.45) |
| Gleason score 7 | 324 (37.63) |
| Gleason score 8 | 53 (6.16) |
| Gleason score 9-10 | 41 (4.76) |
| AJCC Prostate Pathologic stage 7$^{th}$ version | |
| Stage I | 20 (2.32) |
| Stage IIa | 81 (9.41) |
| Stage IIb | 435 (50.52) |
| Stage III | 134 (15.56) |
| Stage IV | 191 (22.18) |
| Prostate Cancer-specific death status, n (%) | |
| No | 835 (96.98) |
| Yes | 26 (3.02) |
| Follow-up duration for prostate cancer survivors in months, median (IQR) | 167 (137–195) |
| Time to cancer-specific death in months, median (IQR) | 116 (73–151) |
| Whole slide imaging | |
| One slide per patient, n (%) | 322 (37.40) |
| Two slides per patient, n (%) | 437 (50.75) |
| Three slides per patient, n (%) | 102 (11.85) |
| Total whole slide images, n | 1,502 |
| Patches with prostate cancer per slide, median (IQR) | 300 (78–735) |
| Total patches with prostate cancer | 847,474 |

### Supplementary table S4

*Table S4 shows the results of comparative analyses between our model, classical and state-of-the-art models that are used in medical imaging. We provided the grade groups as reference. Our novel model delivered a higher generalized concordance probability, a larger effect size or z values than EfficientNet B1, VGG-16 and ResNet-50RS, although our model is computationally efficient (1/3 of EfficientNet B1, 1/76 of VGG-16 or 1/22 of ResNet-50RS), it used fewer features in the final convolutional layer, the last layer before the full connection of these features for prediction (1/20 of EfficientNet B1, 1/8 of VGG-16 or 1/32 of ResNet-50RS) and lesser parameter capacity (1/24 of EfficientNet B1, 1/54 of VGG-16 or 1/125 of ResNet-50RS) to compete with these existing models. We used the univariate weighted cox regression model to estimate generalized concordance probability, the effect size, the z value, the Akaike information criterion (AIC) and the Bayesian information criterion (BIC). The best measured metrics are highlighted in bold. The non-nested partial likelihood ratio test for cox regression models revealed that EfficientNet B1 does not fit better than our novel model (z=0.396 p = 0.346) by ΔAIC = ΔBIC = 5.594. The computation efficacy was measured using gFLOPS (giga floating point operations per second) on the input dimension (16×512×512×3 pixels). The lower gFLOPS, the more efficient is the model.*

| | Grade groups | our novel model | EfficientNet B1 | VGG-16 | ResNet50RS |
|---|---|---|---|---|---|
| Parameter capacity | - | **270,357** | 6,576,520 | 14,715,201 | 33,698,337 |
| Number of feature maps in the last convolutional layer | - | **64** | 1,280 | 512 | 2,048 |
| gFLOPS | - | **33.90** | 98.55 | 2567.35 | 746.59 |
| Concordance index | 0.682±0.017 | 0.682±0.018 | **0.688±0.018** | 0.635±0.011 | 0.667±0.018 |
| Generalized concordance probability | 0.875 (0.771–0.936) | **0.927 (0.891–0.952)** | 0.875 (0.837–0.905) | 0.807 (0.748–0.854) | 0.861 (0.818–0.896) |
| Effect size | 7.000 (3.371–14.540) | **12.703 (8.178–19.729)** | 6.995 (5.122–9.553) | 4.171 (2.966–5.865) | 6.205 (0.484–8.586) |
| *z* | 5.218843 | 11.31463 | **12.23275** | 8.213045 | 11.01189 |
| AIC | 3365.577 | 3328.96 | 3323.366 | 3378.062 | 3351.338 |
| BIC | 3369.168 | 3332.551 | 3326.957 | 3381.653 | 3354.929 |

### Supplementary table S5

*Table S5 shows the results of comparative analyses for different objective magnification (10x, 20x and 40x), as well as the Cox deep model concept and the attention aggregation layer (as alternative for average pooling). These results demonstrate that considering objective magnification higher than 10x, attention aggregation layer instead of average pooling, as well as the cox deep model concept resulted in inferior performance. We used the univariate weighted cox regression model to estimate generalized concordance probability, the effect size, the z value, the Akaike information criterion (AIC) and the Bayesian information criterion (BIC).*

| | Our strategy | Alternative strategies | | | |
|---|---|---|---|---|---|
| | 10x objective magnification | 20x objective magnification | 40x objective magnification | Attention aggregation layer | Cox deep model |
| Concordance index | **0.682±0.018** | 0.660±0.018 | 0.557±0.018 | 0.610±0.011 | 0.588±0.018 |
| Generalized concordance probability | **0.927 (0.891–0.952)** | 0.861 (0.818–0.896) | n.c. | 0.838 (0.756–0.896) | 0.719 (0.627–0.797) |
| Effect size | **12.703 (8.178–19.729)** | 6.205 (4.484–8.586) | n.c. | 5.173 (3.095–8.646) | 2.563(1.677–3.916) |
| *z* | **11.31463** | 11.01189 | 3.201225 | 6.27067 | 4.35174 |
| AIC | **3328.960** | 3387.361 | 3460.59 | 3428.935 | 3446.709 |
| BIC | **3332.551** | 3390.952 | 3464.181 | 3432.526 | 3450.300 |

### Supplementary table S6

*Table S6: Univariate and multivariate weighted Cox regression analyses for recurrence (BCR) in the first validation set (CPCBN, Canada). Age at diagnosis and positive surgical margins (PSM) were not significant nor independent predictors for BCR. Therefore, the multivariate analysis excluded both parameters. CI: confidence interval. z was only reported in the multivariate analysis. PSM: positive surgical margin (reference: negative). PSA: Prostate-specific antigen. IDC: Presence of intraductal carcinoma of the prostate (reference: absent). *Log- transformed.*

| | Univariate analysis | | Multivariate analysis | | | |
|---|---|---|---|---|---|---|
| Parameter | Hazard ratio (95% CI) | *p* | Hazard ratio (95% CI) | *z* | *p* | Generalized concordance probability (95% CI) |
| Age at diagnosis | 1.01 (0.99–1.03) | 0.1578 | - | - | - | - |
| PSA level* | 1.91 (1.50–2.44) | <0.0001 | 1.41 (1.12–1.76) | 2.92 | 0.0345 | 0.505 (0.503–0.507) |
| IDC | 1.83 (1.22–2.76) | 0.0036 | 1.28 (0.79–2.09) | 1.00 | 0.3150 | 0.566 (0.452–0.674) |
| PSM | 0.75 (0.11–5.05) | 0.7631 | - | - | - | - |
| Tumor stage | | | | | | |
| pT2 | Reference | | Reference | | | |
| pT3a | 2.43 (1.80–3.29) | <0.0001 | 1.75 (1.27–2.43) | 3.39 | 0.0071 | 0.643 (0.566–0.713) |
| pT3b | 5.90 (4.21–8.26) | <0.0001 | 3.03 (2.07–4.43) | 5.72 | <0.0001 | 0.754 (0.676–0.818) |
| pT4 | 6.74 (3.41–13.30) | <0.0001 | 3.31 (1.29–8.52) | 2.49 | 0.0128 | 0.750 (0.525–0.891) |
| Grade groups | | | | | | |
| GG1 | Reference | | Reference | | | |
| GG2 | 1.91 (1.41–2.59) | <0.0001 | 1.37 (0.92–2.04) | 1.55 | 0.1202 | 0.591 (0.493–0.682) |
| GG3 | 2.54 (4.36–8.71) | <0.0001 | 1.49 (0.95–2.34) | 1.72 | 0.0848 | 0.614 (0.502–0.714) |
| GG4 | 4.96 (2.89–8.49) | <0.0001 | 1.52 (0.88–2.63) | 1.51 | 0.1308 | 0.613 (0.479–0.731) |
| GG5 | 6.92 (3.16–15.15) | <0.0001 | 2.54 (1.47–4.42) | 3.32 | 0.0090 | 0.731 (0.612–0.825) |
| **BCR score** | **12.70 (8.18–19.73)** | **<0.0001** | **6.43 (3.84–10.77)** | **7.07** | **<0.0001** | **0.867 (0.794–0.917)** |

### Supplementary table S7

*Table S7: multivariate weighted Cox regression analyses of the Grade Groups and the novel risk groups for biochemical recurrence (BCR) on the second validation set (PROCURE, Canada) in addition to preoperative prostate-specific antigen (PSA), tumor staging classification (pT and pN) and positive surgical margin. CI: confidence interval. The hazard ratios for BCR are calculated. CI: confidence interval. *Log- transformed.*

| Parameter | Hazard ratio (95% CI) | *z* | *p* | Generalized concordance probability (95% CI) |
|---|---|---|---|---|
| PSA level* | 1.55 (1.25–1.93) | 4.03 | <0.0001 | 0.609 (0.556–0.658) |
| Tumor stage | | | | |
| pT2 | Reference | | | |
| pT3a | 2.00(1.48–2.71) | 4.46 | <0.0001 | 0.667 (0.596–0.731) |
| pT3b | 3.23 (2.13–4.89) | 5.53 | <0.0001 | 0.764 (0.681–0.830) |
| Nodal stage | | | | |
| pN1 vs. pN0/x | 1.48 (0.83–2.64) | 1.34 | 0.1818 | 0.597 (0.454–0.725) |
| Surgical margin status | | | | |
| Positive vs. Negative | 2.29 (1.69–3.09) | 5.38 | <0.0001 | 0.696 (0.629–0.756) |
| Grade Groups | | | | |
| GG1 | Reference | | | |
| GG2 | 2.45 (1.18–5.13) | 2.82 | 0.0169 | 0.711 (0.540–0.837) |
| GG3 | 3.67 (1.74–7.76) | 3.40 | 0.0006 | 0.786 (0.634–0.886) |
| GG4 | 5.15 (2.08–12.77) | 3.54 | 0.0004 | 0.837 (0.675–0.927) |
| GG5 | 8.12 (3.54–18.63) | 4.94 | <0.0001 | 0.890 (0.830–0.959) |
| BCR score-based risk groups | | | | |
| Low | Reference | | | |
| Low-intermediate | 3.44 (1.99–5.94) | 4.44 | <0.0001 | 0.775 (0.666–0.856) |
| High-intermediate | 6.84 (3.80–12.30) | 6.42 | <0.0001 | 0.872 (0.792–0.925) |
| High | 12.73 (6.91–23.46) | 8.16 | <0.0001 | 0.927 (0.874–0.959) |

#### Supplementary table S8

*Table S8: Biochemical recurrence (BCR)-free survival probabilities for all risk groups based on BCR scores at different time points in the external validation sets (CPCBN and PROCURE). CI: confidence interval.*

| BCR score-based risk groups | BCR-free survival rates | | |
|---|---|---|---|
| | 3rd year (95% CI) | 5th year (95% CI) | 10th year (95% CI) |
| **1st external validation set (CPCBN)** (log-rank $p < 0.0001$) | | | |
| Low | 92.1 (87.5–96.9) | 90.4 (85.3–95.7) | 88.2 (82.5–94.3) |
| Low-intermediate | 86.8 (83.8–90.0) | 83.1 (79.8–86.7) | 75.6 (71.3–80.2) |
| High-intermediate | 78.1 (72.2–84.4) | 69.7 (63.2–76.9) | 54.0 (45.6–63.8) |
| High | 54.0 (45.5–64.0) | 36.7 (28.8–46.9) | 23.7 (16.1–35.1) |
| **2nd external validation set (PROCURE)** (log-rank $p < 0.0001$) | | | |
| Low | 94.0 (90.6–97.5) | 91.6 (87.6–95.8) | 85.7 (79.2–92.8) |
| Low-intermediate | 79.2 (75.8–82.7) | 71.7 (67.9–75.7) | 64.4 (60.1–69.0) |
| High-intermediate | 44.9 (37.3–54.2) | 37.3 (29.8–46.6) | 28.5 (21.0–38.7) |
| High | 19.4 (9.4–39.7) | 9.7 (3.3–28.4) | - |

#### Supplementary table S9

*Table S9: recurrence-free survival rates for all grade groups based at different time points in the first external validation set (CPCBN, Canada). CI: confidence interval.*

| Grade Groups | Recurrence cases | Recurrence-free survival rates (log rank p<0.0001) | | |
|---|---|---|---|---|
| | n (%) | 3rd year (95% CI) | 5th year (95% CI) | 10th year (95% CI) |
| GG1 | 40 (14.8) | 93.3 (90.3–96.3) | 88.9 (85.1–92.8) | 83.0 (77.9–88.4) |
| GG2 | 90 (26.1) | 85.9 (82.2–89.6) | 79.9 (75.7–84.3) | 68.6 (62.8–74.9) |
| GG3 | 61 (41.5) | 73.9 (67.1–81.4) | 68.8 (61.7–76.8) | 53.3 (44.6–63.6) |
| GG4 | 45 (56.9) | 58.0 (48.1–70.0) | 45.8 (35.9–58.5) | 42.2 (32.2–55.3) |
| GG5 | 32 (66.7) | 47.9 (35.7–64.4) | 37.5 (26.0–54.0) | 17.3 (10.1–29.8) |

### Supplementary table S10

*Table S10: recurrence-free survival rates for all grade groups based at different time points in the second external validation set (PROCURE, Canada). CI: confidence interval.*

| Grade Groups | Recurrence cases | Recurrence-free survival rates (log rank p<0.0001) | | |
|---|---|---|---|---|
| | n (%) | 3rd year (95% CI) | 5th year (95% CI) | 10th year (95% CI) |
| GG1 | 15 (10.5) | 95.7 (92.5–99.1) | 94.2 (90.3–98.2) | 88.5 (82.8–94.4) |
| GG2 | 130 (28.6) | 81.9 (78.4–85.6) | 75.9 (71.9–80.1) | 62.9 (56.6–70.0) |
| GG3 | 111 (54.4) | 64.7 (58.4–71.6) | 54.8 (48.2–62.3) | 34.8 (27.2–44.6) |
| GG4 | 17 (63.0) | 48.1 (32.6–71.2) | 37.0 (22.6–60.6) | 37.0 (22.6–60.6) |
| GG5 | 56 (82.4) | 24.5 (16.1–37.3) | 17.3 (10.1–29.8) | - |

### Supplementary table S11

*Table S11: Univariate and multivariate Cox regression analyses for cancer-specific mortality on the first external validation set (CPCBN, Canada). CI: confidence interval. Loco-regional lymph node status was not significant in univariate and multivariate analyses and therefore not presented here. Please note that c-index is not suitable for Cox model comparison.*

| | Univariate analysis | | | | Multivariate analysis | | | |
|---|---|---|---|---|---|---|---|---|
| Parameter | Hazard ratio (95% CI) | *p* | c-index | AIC (BIC) | Hazard ratio (95% CI) | *z* | *p* | c-index |
| Tumor stage | 2.58 (1.58–4.21) | 0.0001 | 0.788 | 171.0623 (171.7703) | 1.89 (1.07–3.34) | 2.20 | 0.0275 | 0.850 |
| Grade Groups | 1.92 (1.30–2.84) | 0.0010 | 0.670 | 173.2988 (174.0069) | 1.33 (0.85–2.09) | 1.24 | 0.2164 | |
| Risk score | 2.58 (1.47–4.55) | 0.0010 | 0.767 | 172.3737 (173.0817) | 1.83 (1.002–3.35) | 1.97 | 0.0491 | |

### Supplementary table S12

*Table S12: univariate and multivariate weighted Cox regression analyses for tumor and nodal stages and the risk scores (continuous parameter) on the second external validation set (PROCURE, Canada). The hazard ratios for cancer-specific mortality are calculated. Due to the limited cancer-specific death events, we binarized the tumor stage (pT2 vs. pT3). The c-index for multivariate analysis is 0.921. CI: confidence interval. Please note that c-index is not suitable for Cox model comparison.*

| | Univariate analysis | | | | Multivariate analysis+ | | | |
|---|---|---|---|---|---|---|---|---|
| Parameter | Hazard ratio (95% CI) | *p* | c-index | AIC (BIC) | Hazard ratio (95% CI) | *z* | *p* | Generalized concordance probability |
| Tumor stage | | | | | | | | |
| pT3 vs. pT2 | 1.97 (0.32 – 12.17) | 0.4632 | 0.744 | 207.54 (208.58) | 0.30 (0.02–4.10) | -0.91 | 0.3654 | 0.230 (0.021–0.804) |
| Nodal stage | | | | | | | | |
| pN1 vs. pN0/x | 21.43 (6.80 – 67.56) | <0.0001 | 0.622 | 202.32 (203.36) | 8.25 (1.36–49.91) | 2.30 | 0.0216 | 0.892 (0.577–0.980) |
| Grade Groups | 76.06 (9.18 – 630.43) | <0.0001 | 0.866 | 183.06 (184.11) | 12.84 (0.87 – 189.84) | 1.86 | 0.0630 | 0.928 (0.465–0.995) |
| Risk score | 109.06 (26.42 – 450.16) | <0.0001 | 0.867 | 185.06 (186.11) | 64.69 (2.20 – 898.43) | 2.42 | 0.0156 | 0.985 (0.688–1.000) |

### Supplementary table S13

*Table S13: univariate and multivariate Cox regression analyses for cancer-specific mortality on the third external validation set (PLCO, U.S.). Note: These Gleason scores were determined prior the introduction of the 2014/2016 ISUP Gleason grading.*

| | Univariate analysis | | | | Multivariate analysis | | | |
|---|---|---|---|---|---|---|---|---|
| Parameter | Hazard ratio (95% CI) | *p* | c-index | AIC (BIC) | Hazard ratio (95% CI) | *Z* | *p* | Generalized concordance probability |
| Gleason score | 54.83 (18.35–163.83) | <0.0001 | 0.806 | 304.5 (305.8) | 34.77 (12.40–97.52) | 6.744 | <0.0001 | 0.972 (0.925–0.990) |
| Risk score | 33.36 (8.70–127.94) | <0.0001 | 0.738 | 315.8 (317.1) | 16.17 (4.60–56.81) | 4.341 | <0.0001 | 0.942 (0.822–0.983) |

### Supplementary table S14

*Table S14 reveals the results of Fine-Gray competing risk regression model for cancer-specific mortality (sub-distribution hazard ratios) on the first external validation set (CPCBN, Canada).* HR reflects the ordering of the risk groups. The extreme high HRs for the risk groups are due to absent of cancer-specific death cases in the low-risk group. The number of death cases from causes other than Prostate Cancer was 72.*

| Characteristic | HR | 95% CI | p-value |
|---|---|---|---|
| Grade Groups | | | |
| GG1 | — | — | |
| GG2 | 0.33 | (0.06 – 2.01) | 0.2 |
| GG3 | 0.56 | (0.09 – 3.42) | 0.5 |
| GG4 | 0.41 | (0.06 – 2.82) | 0.4 |
| GG5 | 2.01 | (0.31 – 13.0) | 0.5 |
| Tumor stage | | | |
| T2 | — | — | |
| T3a | 4.94 | (0.92 – 26.5) | 0.063 |
| T3b | 10.4 | (1.56 – 68.9) | 0.015 |
| T4 | 5.28 | (0.27 – 104) | 0.3 |
| The risk groups* | | | |
| Low | — | — | |
| Low-intermediate | 11,850 | (4,342 – 32,343) | <0.001 |
| High-intermediate | 10,871 | (2,875 – 41,103) | <0.001 |
| High | 32,320 | (9,682 – 107,892) | <0.001 |
| HR = Hazard Ratio, CI = Confidence Interval | | | |

### Supplementary table S15

*Table S15 reveals the results of Fine-Gray competing risk regression model for cancer-specific mortality (sub-distribution hazard ratios) on the second external validation set (PROCURE, Canada).* HR reflects the ordering of the risk groups. The extreme high HRs  for the risk groups and Grade groups are due to absent of cancer-specific death cases in the low-risk group and GG1. The number of death cases from causes other than Prostate Cancer was 69.*

| **Characteristic** | **HR** | **95% CI** | **p-value** |
|---|---|---|---|
| Grade Groups | | | |
| GG1 | — | — | |
| GG2 | 6,111 | (2,630 – 14,200) | <0.001 |
| GG3 | 3,917 | (771 – 19,903) | <0.001 |
| GG4 | 1,470 | (29.4 – 73,470) | <0.001 |
| GG5 | 11,532 | (1,413 – 94,142) | <0.001 |
| Tumor stage | | | |
| T2 | — | — | |
| T3a | 0.82 | (0.18 – 3.71) | 0.8 |
| T3b | 2.60 | (0.71 – 9.55) | 0.2 |
| Lymph node stage | | | |
| pN0/x | — | — | |
| pN1 | 0.40 | (0.13 – 1.26) | 0.12 |
| The risk groups* | | | |
| Low | — | — | |
| Low-intermediate | 6,797 | (3,002 – 15,390) | <0.001 |
| High-intermediate | 17,425 | (3,549 – 85,551) | <0.001 |
| High | 106,639 | (18,456 – 616,165) | <0.001 |
| HR = Hazard Ratio, CI = Confidence Interval | | | |

### Supplementary table S16

*Table S16 reveals the results of Fine-Gray competing risk regression model for cancer-specific mortality (sub-distribution hazard ratios) on the third external validation set (PLCO, US). The extreme high HRs for the risk groups and pathologic Prostate Stage are due to absent of cancer-specific death cases in the low-risk group and Stage I. The number of death cases from causes other than Prostate Cancer was 179.*

| **Characteristic** | **HR** | **95% CI** | **p-value** |
|---|---|---|---|
| Gleason score | | | |
| 6 | — | — | |
| 7 | 5.92 | (1.27 – 27.5) | 0.023 |
| 8 – 9 | 22.1 | (3.63 – 134) | <0.001 |
| 10 | 16.5 | (2.48 – 110) | 0.004 |
| Pathologic Prostate Stage | | | |
| Stage I | — | — | |
| Stage IIA | 0.80 | (0.15 – 4.12) | 0.8 |
| Stage IIB | 16,481 | (4,634 – 58,614) | <0.001 |
| Stage III | 17,348 | (5,580 – 53,935) | <0.001 |
| Stage IV | 24,404 | (8,783 – 67,808) | <0.001 |
| The novel risk groups* | | | |
| Low | — | — | |
| Low-intermediate | 12,156 | (5,759 – 25,656) | <0.001 |
| High-intermediate | 29,687 | (14,108 – 62,469) | <0.001 |
| High | 35,406 | (9,912 – 126,466) | <0.001 |
| HR = Hazard Ratio, CI = Confidence Interval | | | |

### Supplementary table S17

*Table S17: Cancer-specific survival rates for all risk groups at different time points in three external validation sets. $\chi^2$ test shows a significant difference for the cancer-specific death status between the risk groups. The low-risk group has no cancer-specific death cases, resulting in the most favorable cancer-specific survival rates of 100%. SE, Standard Error. In the PROCURE cohort, estimating 15-year cancer-specific survival rates was not feasible due to the constraint of the follow-up duration.*

| Risk groups | Cancer-specific death cases | Cancer-specific survival rates | |
|---|---|---|---|
| | n (%) | $10^{th}$ year (SE) | $15^{th}$ year (SE) |
| **$1^{st}$ external validation set (CPCBN)** (log-rank p = 0.0014; $\chi^2(3, N=889) = 17.121$, p = 0.0007) | | | |
| Low | 0 (0) | 100.0 (0) | 100.0 (0) |
| Low-intermediate | 5 (1.1) | 99.3 (0.5) | 96.8 (1.5) |
| High-intermediate | 3 (1.7) | 98.2 (1.3) | 96.1 (2.9) |
| High | 7 (6.2) | 92.7 (2.3) | 89.0 (4.6) |
| **$2^{nd}$ external validation set (PROCURE)** (log-rank p < 0.0001; $\chi^2(3, N=897) = 67.063$, p <0.0001) | | | |
| Low | 0 (0) | 100.0 (0) | n.c. |
| Low-intermediate | 7 (1.3) | 96.6 (1.7) | n.c. |
| High-intermediate | 7 (5.1) | 72.8 (9.3) | n.c. |
| High | 7 (22.6) | 60.3 (14.1) | n.c. |
| **$3^{rd}$ external validation set (PLCO)** (log-rank p < 0.0001; $\chi^2(3, N=861) = 19.023$, p = 0.0003) | | | |
| Low | 0 (0) | 100.0 (0) | 100.0 (0) |
| Low-intermediate | 11 (2.0) | 98.9 (0.5) | 97.8 (0.7) |
| High-intermediate | 12 (6.3) | 95.5 (1.5) | 93.0 (2.1) |
| High | 3 (12.5) | 95.7 (4.3) | 81.2 (10.2) |

### Supplementary table S18

*Table S18: Multivariate Cox regression analysis including risk scores and the slide number on the third external validation set (PLCO, U.S.). Hazard ratio for cancer-specific mortality is estimated. The results indicate no significant interaction between the risk score and the slide number; the Pearson correlation coefficient between the risk score and the slide number was 0.005 (95% CI: -0.062–0.072; p=0.8792).*

| Parameter | Hazard ratio (95% CI) | *z* | *p* |
|---|---|---|---|
| Risk score | 52.93 (2.47–1135.56) | 2.537 | 0.0112 |
| Slide number | 1.91 (0.91–4.00) | 1.710 | 0.0872 |
| **Slide number · risk score** | **0.89 (0.19–4.12)** | **-0.146** | **0.8839** |

### Supplementary table S19

*Table S19: cancer-specific survival rates for all grade groups based at different time points in the first external validation set (CPCBN, Canada). SE: Standard Error.*

| Grade Groups | Cancer-specific death cases | Cancer-specific survival rates (log rank p<0.0001) | |
|---|---|---|---|
| | n (%) | 10th year (SE) | 15th year (SE) |
| GG1 | 3 (1.1) | 99.2 (0.6) | 98.0 (1.3) |
| GG2 | 2 (0.6) | 99.3 (0.6) | 98.0 (1.5) |
| GG3 | 3 (2.0) | 98.0 (1.4) | 96.1 (2.3) |
| GG4 | 2 (2.5) | 100 (0) | 92.2 (5.3) |
| GG5 | 5 (10.4) | 83.0 (7.0) | 83.0 (7.0) |

#### Supplementary table S20

*Table S20: cancer-specific survival rates for all grade groups based at different time points in the second external validation set (PROCURE, Canada). SE: Standard Error.*

| Grade Groups | Cancer-specific death cases | Cancer-specific survival rates (log rank p<0.001) |
|---|---|---|
| | n (%) | $10^{th}$ year (SE) |
| GG1 | 0 (100) | 100 (0) |
| GG2 | 6 (1.3) | 95.5 (2.1) |
| GG3 | 4 (2.0) | 96.4 (2.9) |
| GG4 | 1 (3.7) | 93.8 (6.1) |
| GG5 | 10 (14.7) | 45.2 (14.2) |

### Supplementary table S21

*Table S21: Cancer-specific survival rates for Gleason scores at different time points in the third external validation set (PLCO, U.S.). These Gleason scores were determined according to prior grading guidelines because the PLCO patients were staged and graded between 1993 and 2008. SE: Standard Error.*

| Gleason score | Cancer-specific death cases | Cancer-specific survival rates (log rank p<0.0001) | |
|---|---|---|---|
| | n (%) | 10th year (SE) | 15th year (SE) |
| GS 6 | 2 (0.45) | 99.8 (0.24) | 99.4 (0.40) |
| GS 7 | 12 (3.70) | 98.3 (0.74) | 95.6 (1.41) |
| GS 8 | 7 (13.21) | 91.8 (3.96) | 82.4 (6.43) |
| GS 9-10 | 5 (12.20) | 87.9 (5.70) | 80.6 (8.74) |

### Supplementary figure S1

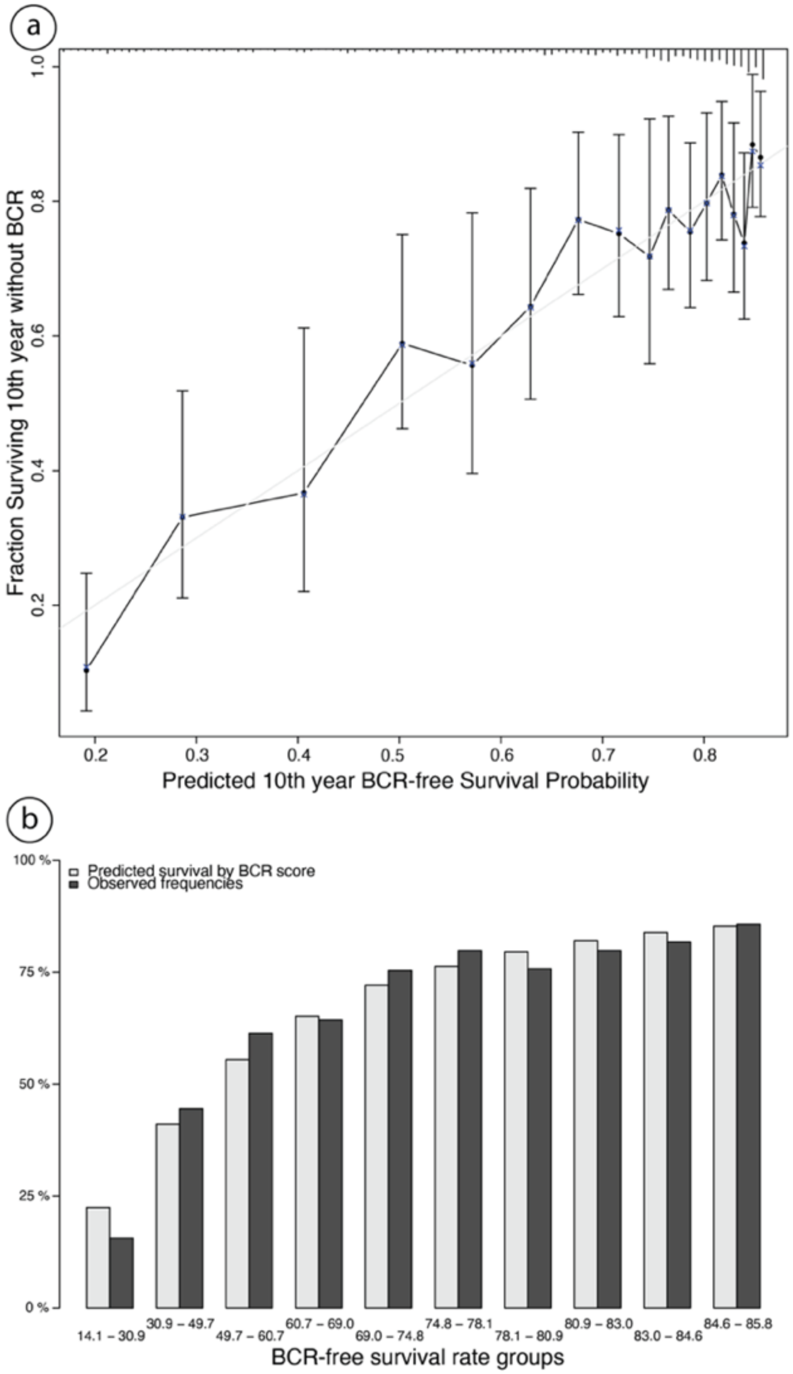

*Figure 1: a. The calibration plot for our biochemical recurrence (BCR)-free prediction model at the 10th year with the rug plot (upper line) for distribution visualization. b. The histogram for observed and predicted survival by BCR scores within groups defined by deciles for observed BCR-free survival rates in the first external Validation set.*

### Supplementary figure S2

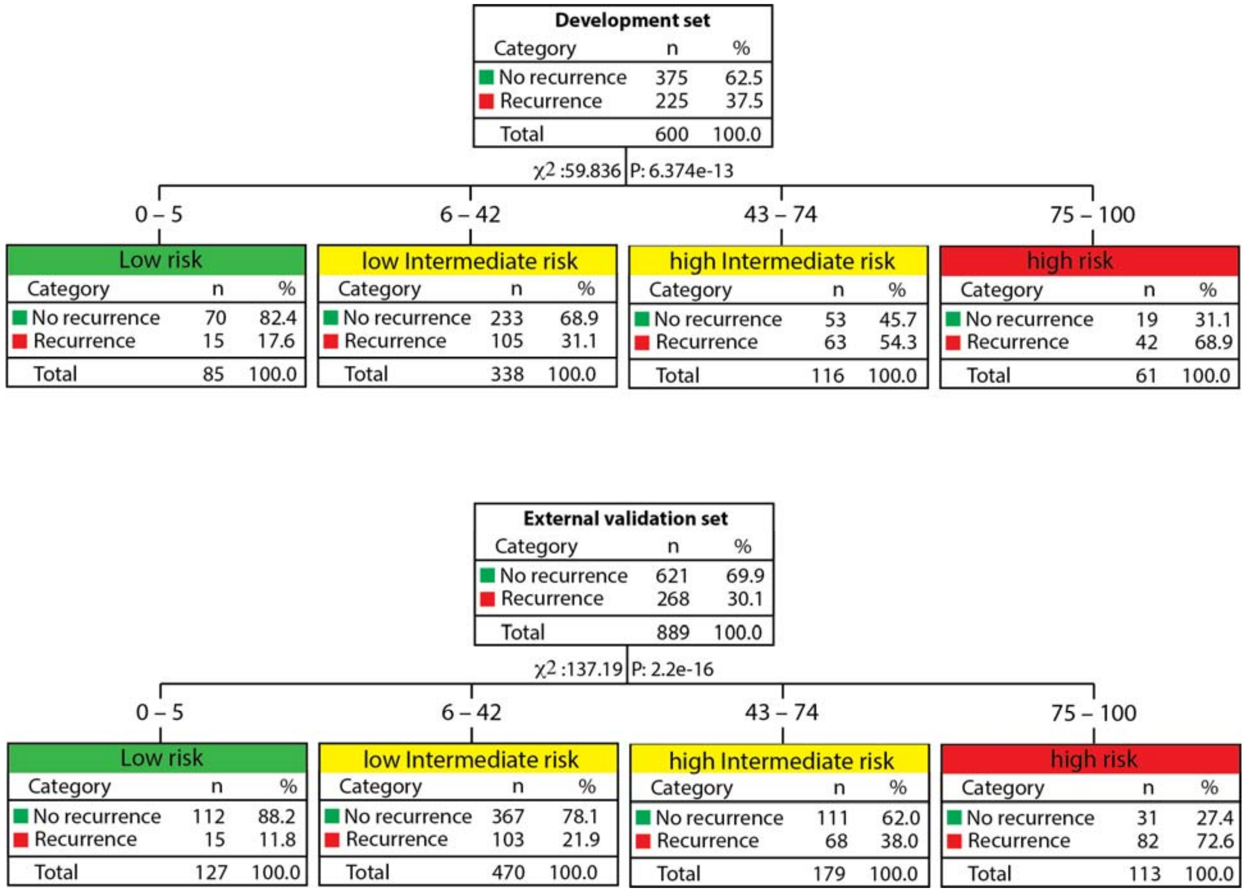

*Figure 2: Risk strata according to the BCR score and corresponding BCR-free rates. Chi-square Automatic Interaction Detection (CHAID) was applied to the Development set to determine the cutoffs for these risk groups. The resulting cutoffs for the BCR scores were fixed for the external Validation set. $X^2$ represents Chi-square statistics.*

### Supplementary figure S3

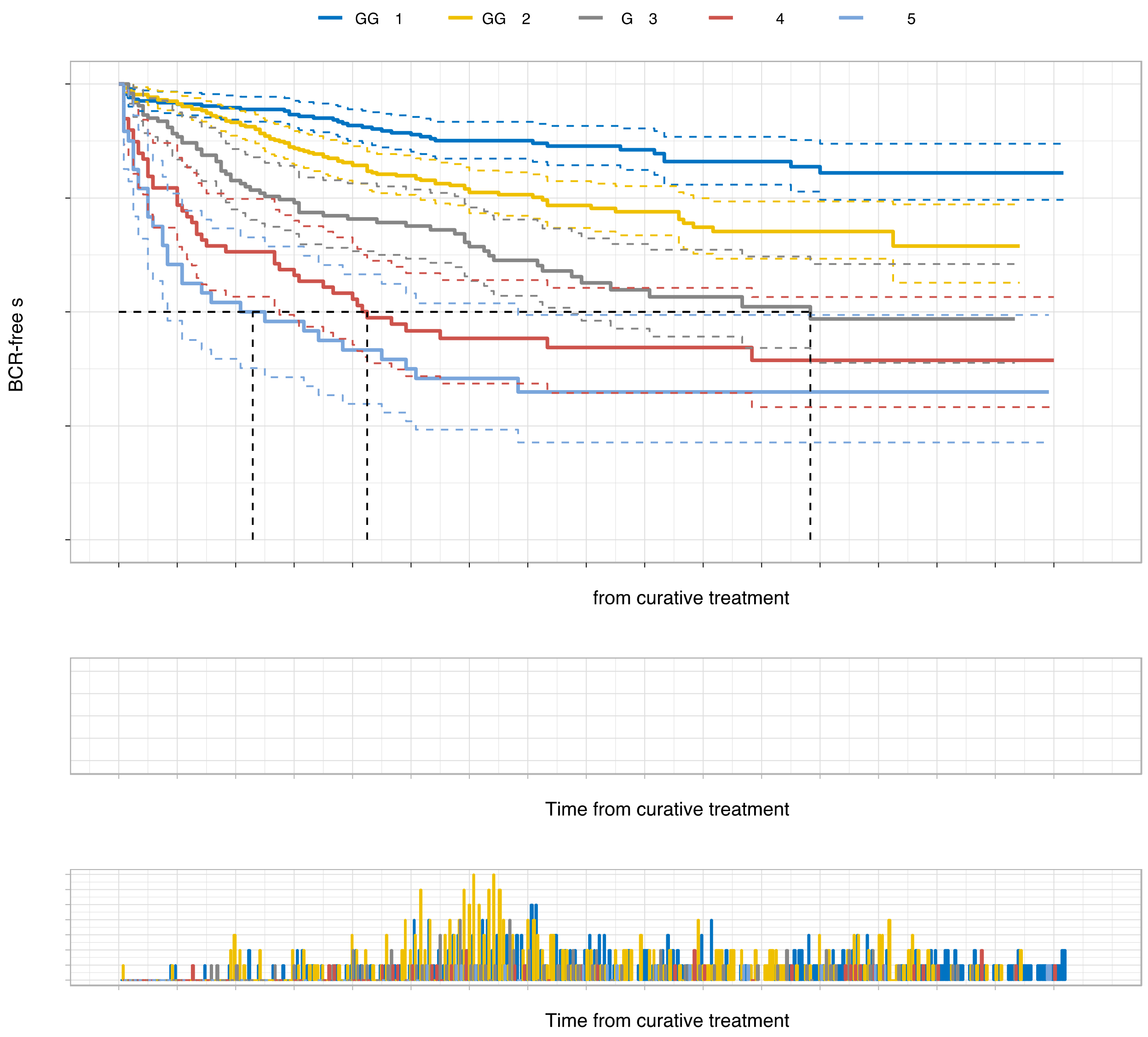

*Figure S3: Kaplan-Meier curves of biochemical recurrence (BCR)-free survival according to grade groups (GG) in the first external validation set (CPCBN, Canada). The dotted lines indicate the median survival. The p-value was measured using the log-rank test. The number of patients at risk and of censored observations are provided for the follow-up period.*

### Supplementary figure S4

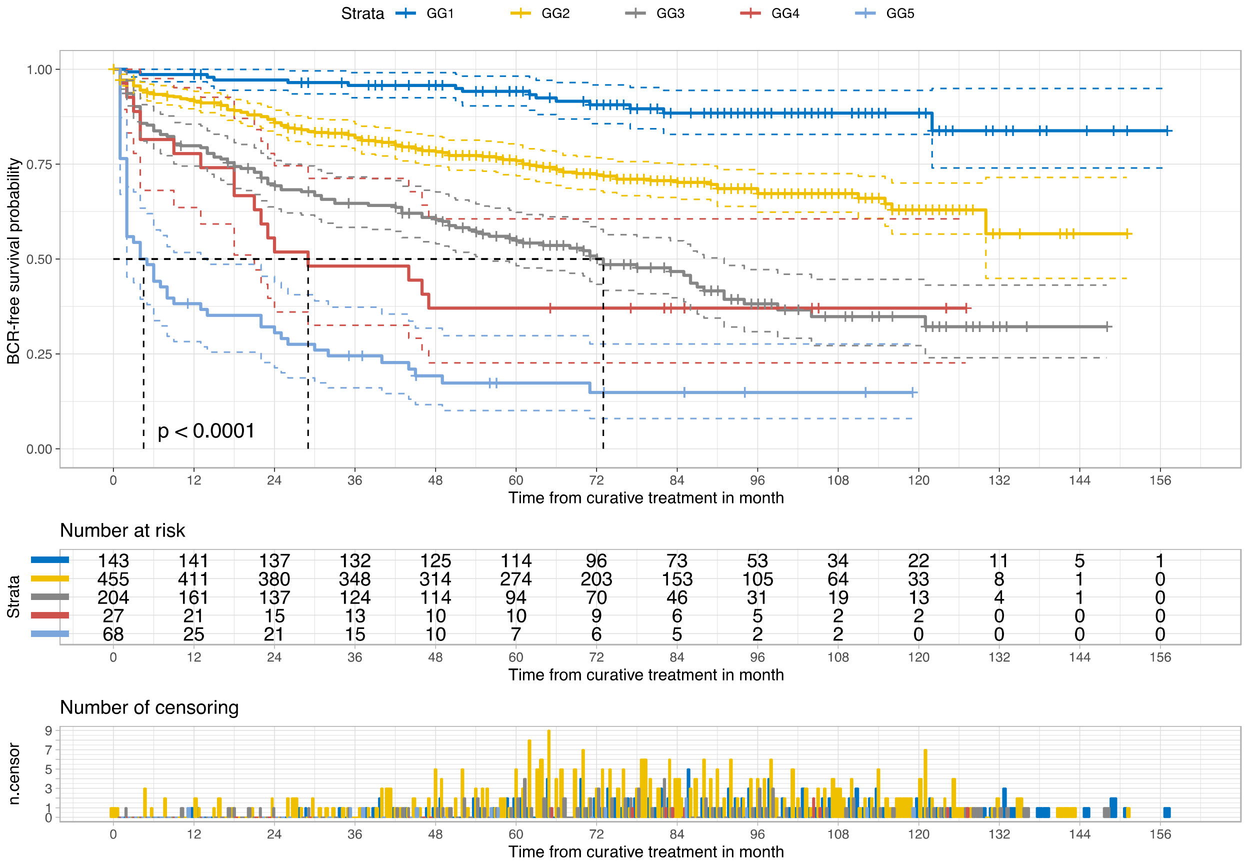

*Figure S4: Kaplan-Meier curves of biochemical recurrence (BCR)-free survival stratified by grade groups (GG) in the second external validation set (PROCURE, Canada). The dotted lines indicate the median survival. The p-value was measured using the log-rank test. The number of patients at risk and of censored observations are provided for the follow-up period.*

### Supplementary figure S5

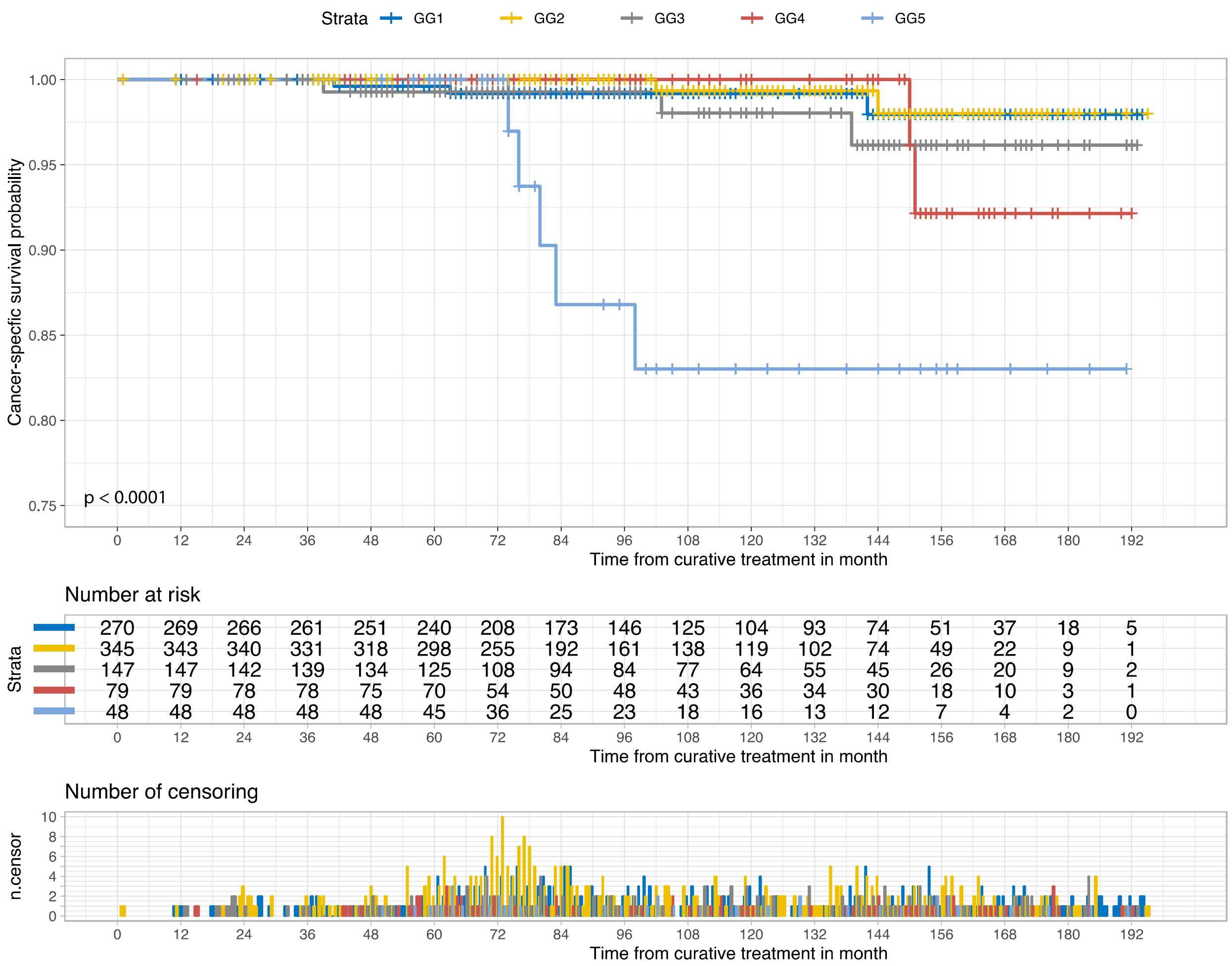

*Figure S5: Kaplan-Meier curves of cancer-specific survival stratified by grade groups (GG) in the first external validation set (CPCBN, Canada)The p-value was measured using the log-rank test. The number of patients at risk and of censored observations are provided for the follow-up period.*

### Supplementary figure S6

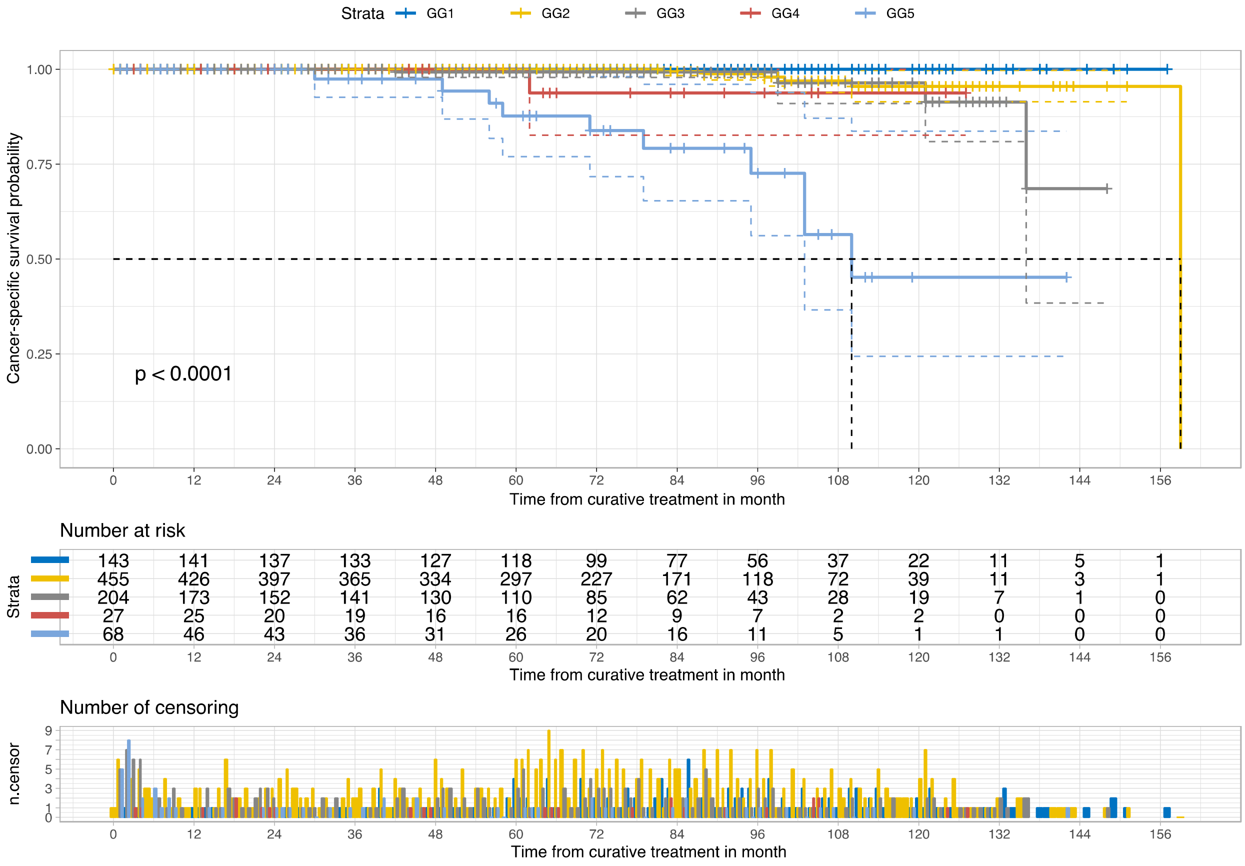

*Figure S6: Kaplan-Meier curves of cancer-specific survival stratified by grade groups (GG) in the second external validation set (PROCURE, Canada). The p-value was measured using the log-rank test. The number of patients at risk and of censored observations are provided for the follow-up period.*

### Supplementary figure S7

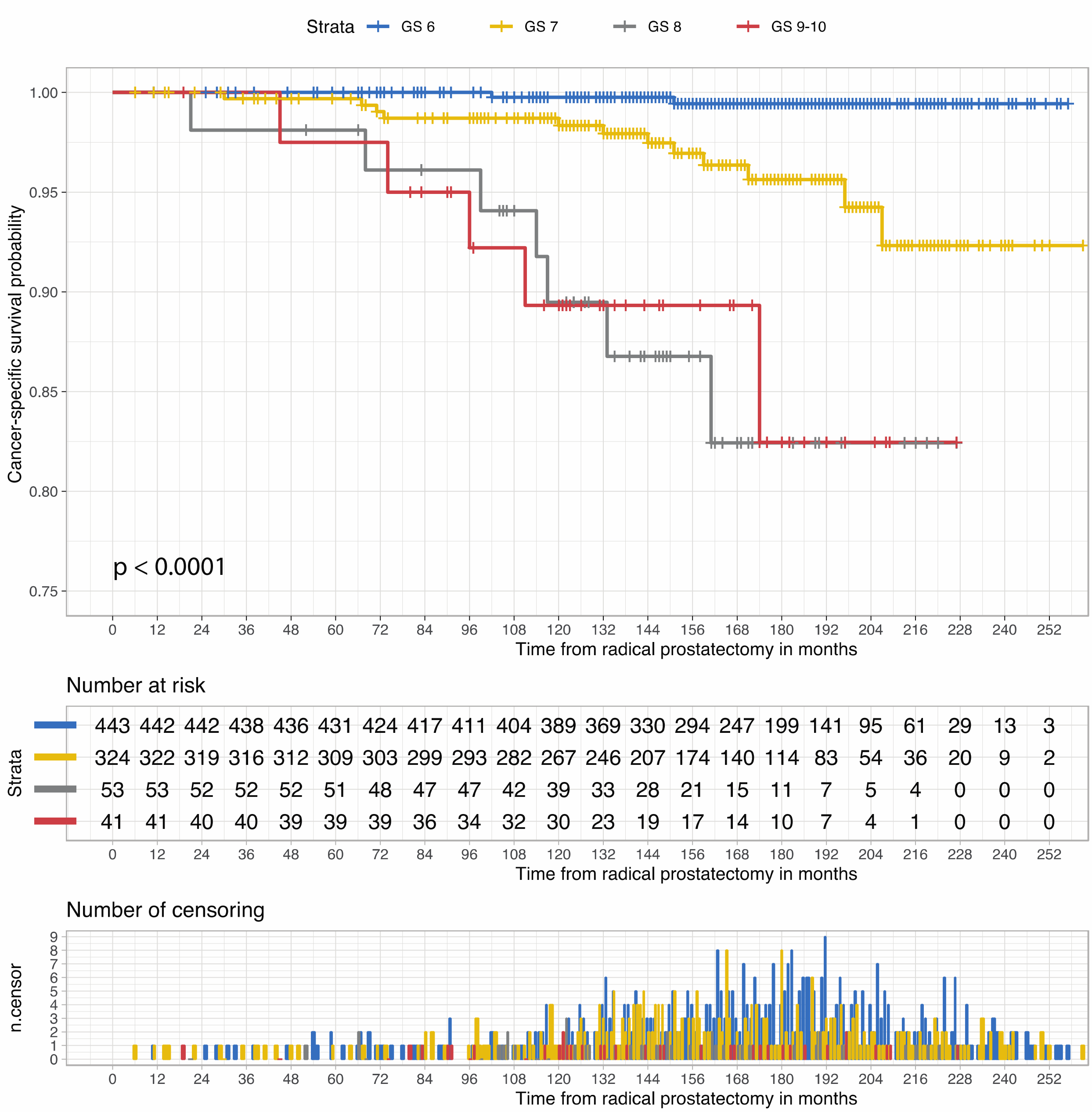

*Figure S7: Kaplan-Meier curves of cancer-specific survival stratified by Gleason scores (GS) in the third external validation set (PLCO, U.S.). The p-value was measured using the log-rank test. The number of patients at risk and of censored observations are provided for the follow-up period.*

### Supplementary figure S8

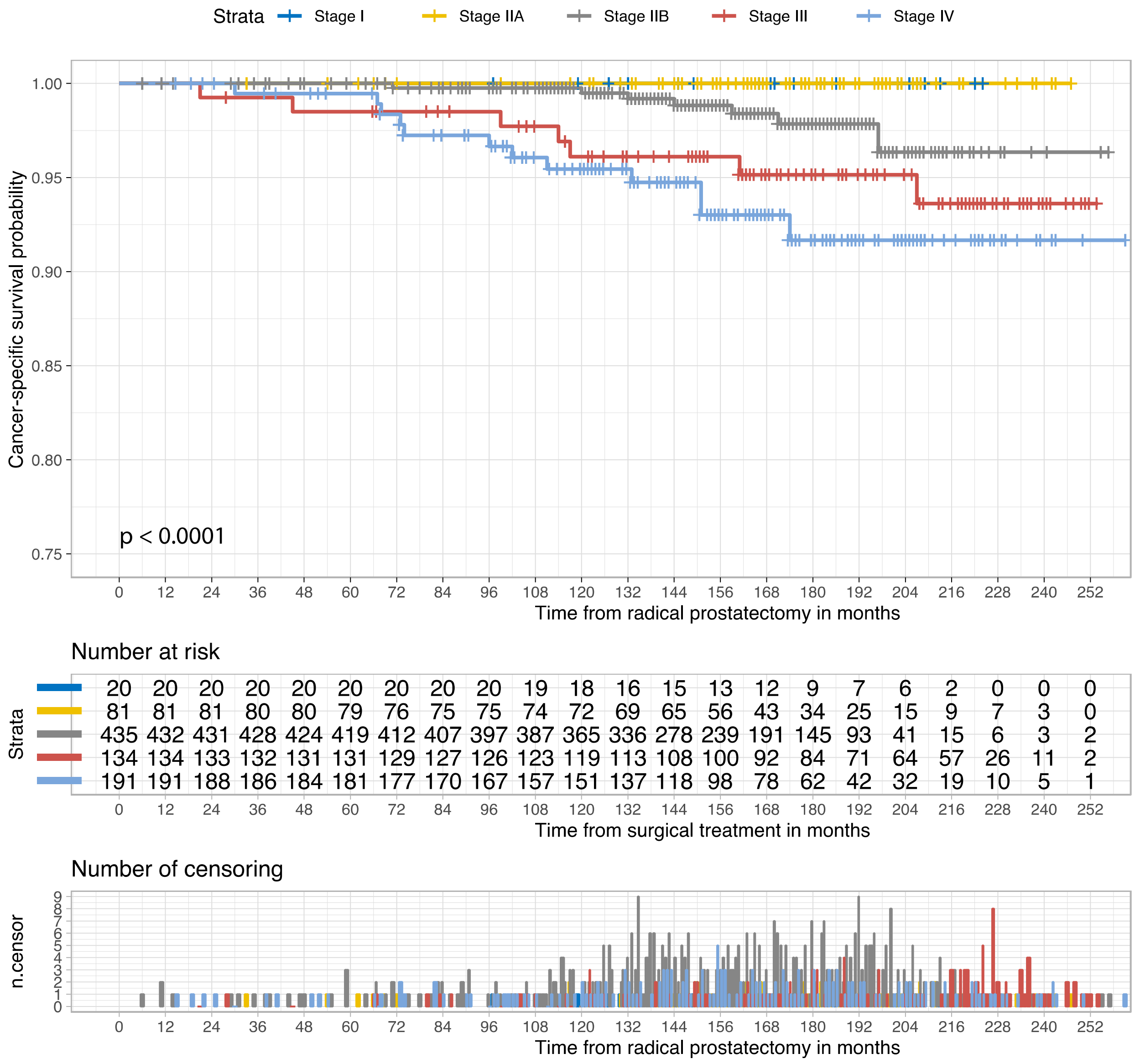

*Figure S8: Kaplan-Meier curves of cancer-specific survival stratified by prostate cancer pathologic stage in the third external validation set (PLCO, U.S.). The p-value was measured using the log-rank test. The number of patients at risk and of censored observations are provided for the follow-up period.*
